# CHD7 binds distinct regions in the *Sox11* locus to regulate neuronal differentiation

**DOI:** 10.1101/2025.04.02.646816

**Authors:** Edward Martinez, Azadeh Jadali, Jingyun Qiu, Anna-Maria Hinman, Julie Zhouli Ni, Jihyun Kim, Kelvin Y. Kwan

**Affiliations:** Department of Cell Biology & Neuroscience, Rutgers University, Piscataway, NJ 08854, USA; Stem Cell Research Center and Keck Center for Collaborative Neuroscience, Rutgers University, Piscataway, NJ 08854, USA; Department of Molecular Biology and Biochemistry, Rutgers University, Piscataway, NJ 08854, USA

## Abstract

The chromodomain helicase DNA binding protein 7 (CHD7) is a nucleosome repositioner implicated in multiple cellular processes, including neuronal differentiation. We identified CHD7 genome-wide binding sites that regulate neuronal differentiation in an otic stem cell line. We identified CHD7 enrichment at the *Sox11* promoter and 3’ untranslated region (UTR). *Sox11* is a transcription factor essential for neuronal differentiation. CRISPRi of *Sox11* promoter or 3’UTR displayed decreased neurite lengths and reduced neuronal marker expression TUBB3 expression. We showed that the *Sox11* locus resides at TAD boundaries, and CTCF marks the 3’UTR. We propose that CHD7 modulates chromatin accessibility of the *Sox11* promoter and CTCF-marked insulators in the 3’UTR to facilitate neuronal differentiation. CRISPRi of the insulator site alters 3D chromatin organization, affects gene expression and ultimately perturbs cellular processes. Our results implicate a general mechanism of CHD7 in facilitating neuronal differentiation and provide insight into CHD7 dysfunction in CHARGE syndrome, a congenital disorder associated with hearing loss.

## Background

Chromatin-modifying enzymes that result in physiological or pathological signals have provided conceptual advances in epigenetic control and changed our understanding of development and disease states (Allis and Jenuwein, 2016). The histone code posits that histone writers, readers, and erasers modify histones that constitute the basis for a gene regulatory code to infer a cell state, identity, and behavior. Many covalent histone modifications have suggested that the nucleosome carries epigenetic information. In addition to readers, writers, and erasers, chromatin regulation involves ATP-dependent nucleosome remodelers, including chromodomain helicase DNA-binding proteins (Clapier et al., 2017). With their enzymatic activities, the CHD family of nucleosome repositioners has significant implications for human health, as pathogenic variants can lead to various diseases, including cancer, neurological disorders, and developmental diseases (Micucci et al., 2015). The CHD proteins form a nine-family-member group of nucleosome repositioners. Deficits in CHD enzymatic activities have provided insight into how altering the cell’s epigenetic landscape affects development and causes disease. Even though there are nine members of CHDs, they each seemingly have distinct cellular functions. Several CHDs and histone deacetylases (HDAC) comprise the catalytic subunits of the nucleosome remodeling (NuRD) complexes (Torrado et al., 2017). NuRD complexes with individual CHD paralogs (CHD3, CHD4, and CHD5) have distinct functions in brain development. CHD4 promotes progenitor proliferation, while CHD5 and CHD3 are involved in neuronal migration and cortical layer specification. Genetic ablation or shRNA knockdown of each CHD leads to unique defects not rescued by the remaining paralogous CHDs (Nitarska et al., 2016).

Understanding each CHD provides insight into maintaining the chromatin landscape for a cell. CHD7 implicates nucleosome remodeling in transcriptional regulation, disease, and embryonic development. Haploinsufficiency of CHD7 causes CHARGE syndrome, a multiple congenital anomaly associated with hearing loss. Pathogenic variants of CHD7 span different domains of the protein in CHARGE syndrome (van Ravenswaaij-Arts and Martin, 2017). In particular, patient mutations from CHD7 show subtle to complete inactivation of chromatin remodeling activity *in vitro*, and links CHD7’s chromatin repositioning activity to CHARGE Syndrome (Bouazoune and Kingston, 2012). A salient feature of CHARGE syndrome is inner ear anomalies. Mice with *Chd7* haploinsufficiency showed conductive and sensorineural hearing loss (Hurd et al., 2011). Conditional knockout and null animals displayed a reduction in the vestibulocochlear ganglion size, a decrease in neuron numbers, and a reduction in proneural gene expression (Hurd et al., 2010). Using a gene-trap reporter mouse suggested that CHD7 is expressed widely in the mature inner and outer hair cells, spiral ganglion neurons (SGNs), vestibular sensory epithelia, and middle ear ossicles, implicating its essential role in different inner ear cell types. CHD7 is expressed in early inner ear neuroblasts and is necessary for the proliferation of inner ear neuroblasts and inner ear morphogenesis (Hurd et al., 2007). In addition to its critical role in inner ear development, CHD7 also has a post-natal role in protecting SGNs and sensory hair cells from stress-induced degeneration (Ahmed et al., 2021). CHD7 likely has multiple developmental and post-natal functions in inner ear cell types.

Linking the developmental process to CHD7’s enrichment at distinct sites in specific cell types is challenging due to the limited number of inner ear cells obtained for interrogating the molecular function of CHD7. Using cellular platforms of inner ear development allows us to understand where CHD7 is binding and how it affects cellular behavior. An inner ear organoid study suggested that it is involved in otic lineage specification and hair cell differentiation (Nie et al., 2022). Genome-wide binding analysis using embryonic stem cells revealed an association of CHD7 with active enhancer marks (Schnetz et al., 2009). CHD7 was associated with cell type-specific enhancers (Reddy et al., 2021; Sanosaka et al., 2022). The studies suggested that CHD7 functions at cell type-specific enhancers to specify the transcriptome of distinct inner ear cell types. Using otic vesicle tissue to identify CHD7 binding, CHD7 binds to promoters and enhancers (Gao et al., 2024) and other undefined cis-regulatory elements. The mechanisms of CHD7 function in the inner ear are not well understood. Cellular systems provide an opportunity to identify novel regulatory regions that require chromatin repositioning activity.

Pluripotent stem cell models offer an approach to generate many cells for molecular studies. Using heterogeneous inner ear cell types is not amenable to probing cell type-specific CHD7 binding. To address this issue, clonally-derived immortalized multipotent otic progenitor (iMOP) cell lines, restricted in cell fate, were developed to generate large numbers of SGN-like cells efficiently. iMOP cells were generated from mouse embryonic (E) 12.5-13.5 cochleae by transient C-MYC expression (Kwan et al., 2015). iMOP cells continue self-renewing in the presence of FGF signaling, and withdrawal of growth factor allows the cells to exit the cell cycle and express cell cycle inhibitors such as CDKN1B (p27^KIP^). Under the appropriate conditions, iMOP cells differentiate into bipolar and pseudounipolar neurons marked by TUBB3 (Tuj-1) (Jadali et al., 2016). Morphologically, the iMOP-derived neurons are reminiscent of SGNs in the cochlea. iMOPs provide a valuable tool for exploring the epigenetic landscape of otic cells during neuronal differentiation.

## Material and Methods

### iMOP cultures

Multipotent otic progenitor cells were grown in suspension with DMEM/F12 (Life Technologies #11320033) containing N21-MAX supplement (R&D Systems #AR008), 25 μg/ml carbenicillin (Fisher Scientific #BP26481), and 20 ng/ml bFGF (PeproTech #450-33). Cells were cultured on a 1.5 coverglass (Electron Microscopy Sciences #72230-01) for immunofluorescence labeling. Coverglass were coated with 10 μg/mL of poly-D-lysine (Corning #354210) for 1 hour, aspirated, and coated overnight with 10 μg/mL of laminin (Life Technologies #23017-015) at 37°C, washed 3 times with 1X PBS before plating cells. For neuronal cultures, cells were dissociated in TrypLE Express (Life Technologies #12604-013) and diluted in neurobasal media before being counted using a Moxi cell counter (Orflo). 0.8 × 10^5^ cells were seeded on single poly-D-lysine and laminin-coated coverslips in a 24-well culture plate and cultured in Neurobasal Medium (Life Technologies #21103049) containing N21-MAX and 2mM L-glutamine (Life Technologies #25030081) and 1μM K03861 (Selleckchem # S8100) at the time of plating. The medium was changed every other day. Immunostaining was performed on 7-day-old cultures.

### Immunohistochemistry

iMOP cells were fixed in 4% formaldehyde in 1X PBS for 20 minutes, incubated in blocking buffer (PBS, 5% goat serum and 0.1% Triton X-100) for 1 hour and incubated overnight with the primary antibody in blocking buffer. Cells were incubated with primary antibodies. After incubation with primary antibodies, cells were rinsed in wash buffer and incubated with appropriate combinations of DAPI (1 µg/ml), Alexa Fluor 488 (1:500 dilution), Alexa Fluor 568 (1:500 dilution) or Alexa Fluor 647 (1:500 dilution) conjugated secondary antibodies (Life Technologies) in blocking buffer for 2 hours. Samples were washed with 1X PBS and mounted in Prolonged Gold Antifade (Thermo Fisher Scientific #P36934). Embryonic tissues were placed in blocking solution (1X PBS with 0.1% Triton X-100 and 5% normal goat serum) for 1 hour at RT and incubated overnight at 4°C with appropriate primary antibodies diluted in blocking solution. After washing with 1X PBS, samples were incubated with combinations of DAPI (1 µg/ml), Alexa Fluor 488 (1:500 dilution), Alexa Fluor 568 (1:500 dilution) or Alexa Fluor 647 (1:500 dilution) conjugated secondary antibodies (Life Technologies) for 2 hours at RT. Samples were washed with 1X PBS and mounted in Prolonged Gold Antifade. Antibodies used are listed in Table 1.

**Table 1:** Antibodies for immunostaining, CUT&Tag and ChIP-seq.

| **Antibody** | **Company** | **Identifier** | **Dilution** | **Purpose** |
| --- | --- | --- | --- | --- |
| Rabbit anti-CHD7 D345 | Cell Signaling | Cat# 6505, RRID:AB_11220431 | 1:300<br>1:50 | Immunostaining<br>CUT&Tag/ChIP-Seq |
| Mouse anti-Tuj1 (TUBB3) | BioLegend | Cat# 801202, RRID:AB_10063408 | 1:300 | Immunostaining |
| Rabbit anti-RFP | Rockland Immunochemicals | Cat# 600-401-379, RRID:AB_2209751 | 1:500 | Immunostaining |
| Goat anti-rabbit Alexa Fluor 488 | Thermo Fisher Scientific | Cat# A-11034, RRID:AB_2576217 | 1:500 | Immunostaining |
| Goat anti-mouse Alexa Fluor 568 | Thermo Fisher Scientific | Cat# A-11004, RRID:AB_2534072 | 1:500 | Immunostaining |
| Goat anti-rabbit Alexa Fluor 568 | Thermo Fisher Scientific | Cat# A-11011, RRID:AB_143157 | 1:500 | Immunostaining |
| Rabbit anti-H3K4me3 | Active Motif | Cat# 39060, RRID:AB_2615077 | 1:50 | CUT&Tag |
| Rabbit anti-EP300 | Santa Cruz Biotechnology | Cat# sc-585, RRID:AB_2231120 | 1:50 | CUT&Tag |
| Rabbit anti-CTCF | Active motif | Cat# 61311, RRID:AB_2614975 | 1:50 | CUT&Tag |
| Rabbit anti-H3K36me3 | Active motif | Cat# 61102, RRID:AB_2615073 | 1:50 | CUT&Tag |
| rabbit IgG | Jackson ImmunoResearch | Cat# 011-000-003, RRID:AB_2337118 | 1:50 | CUT&Tag |

### Cell Proliferation EdU Assay

iMOP cells were plated and cultured in suspension with DMEM/F12 containing N21-MAX supplement, 25 µg/ml carbenicillin, and 20 ng/ml bFGF. Three days after plating, cells were dissociated with TrypLE Express, harvested by centrifugation, and resuspended in culture medium. Cell numbers were counted using a Moxi counter. Particles between 9-13 um were considered cells. 1.5 × 10^5^ cells were plated in poly-D-lysine and laminin-coated coverslips. EdU (final concentration: 10 µM) was added three hours after plating for two hours. Cells were fixed in 4% formaldehyde in 1X PBS for 15 minutes and permeabilized in 0.5% TritonX-100 in 1x PBS buffer for 20 minutes. The Click-iT EdU Imaging kit (Invitrogen #C10337) was used to detect EdU according to the manufacturer’s instructions.

### Transduction, transfection and generation of stable iMOP cell lines

Single-stranded complementary oligos containing either scrambled or *Chd7* shRNA sequence was 5’ phosphorylated with polynucleotide kinase and annealed to form a double-stranded oligo containing restriction site overhangs for cloning into lentiviral vectors. Double-stranded oligos containing the shRNA sequences were ligated lentiviral vectors containing a blasticidin resistance cassette, pLKO.1-blast (Addgene #26655) into the AgeI and EcoRI restriction sites. Viral constructs and packaging plasmids were transfected into 293FT cell line, and the supernatant was collected 48- and 64-hours post-transfection and combined. Viruses were concentrated by precipitation using PEG 6000 (Kutner et al., 2009). For *Chd7* knockdown and detection by Western blot, iMOP cells were infected at a multiplicity of infection (MOI) of 5. Twenty-four hours after infection, cells were selected with 10 µg/ml blasticidin (Thermofisher # A1113903). Cells were harvested 72 hours after selection. For transient shRNA knockdown experiments, iMOP cells were plated for neuronal differentiation and allowed to recover for up to 24 hours before transfection with pLKO.3G *Chd7* shRNA or pLKO.3G scrambled shRNA using jetPRIME (Polyplus # 101000046).

To generate a stable CRISPRi (dCas9-KRAB-MeCP2) cell line, the Lenti_dCas9-KRAB-MeCP2 (Addgene #122205) was used as a lentiviral expression vector. The Lenti_dCas9-KRAB-MeCP2 plasmid with pRSV-Rev (Addgene #12253), pMDLg/pRRE (Addgene #12251) and pMD2.G (VSVG) (Addgene #12259) were used for producing lentiviral particles. HEK293FT packaging cells were transfected with the three aforementioned plasmids using jetOPTIMUS transfection reagent (Polyplus). The supernatant was collected 48- and 72 hours post-transfection. Proliferating iMOPs were infected with unconcentrated lentivirus. Forty-eight hours after infection, cells underwent stepwise blasticidin selection. Cells were eventually selected in 2 µg/ml of blasticidin for 3 weeks to obtain stable CRISPRi cell populations.

sgRNAs that target specific regions of *Sox11* were designed using CRISPick (Doench et al., 2016; Sanson et al., 2018) and the sequences for sgRNA are provided in Table 3. The sgRNA was cloned into a lentiviral vector containing a neomycin resistance cassette by Gibson Assembly (Addgene #41824) and transduced into cells. Cells were selected with neomycin and differentiated into iMOP-derived neurons. iMOP cells were infected at a multiplicity of infection (MOI) of 5. Twenty-four hours after infection, cells were selected by a stepwise increase of the selection drug up to 200 µg/ml G418 (Thermofisher # 11811031). Cells were harvested 72 hours after selection.

**Table 2:**
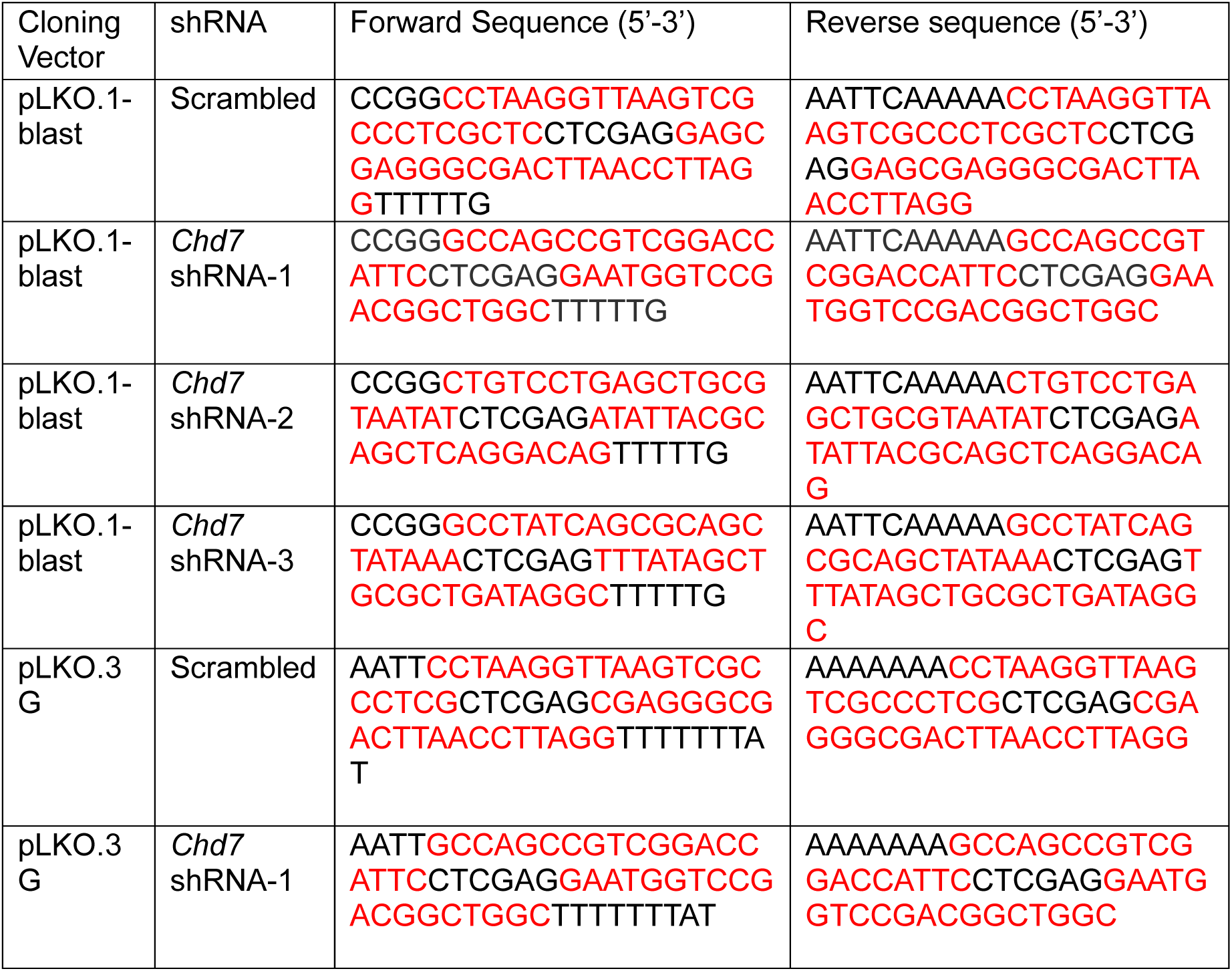
*Chd7* shRNA sequence cloned into lentiviral vectors. Red nucleotides represent the shRNA sequence.

**Table 3:**
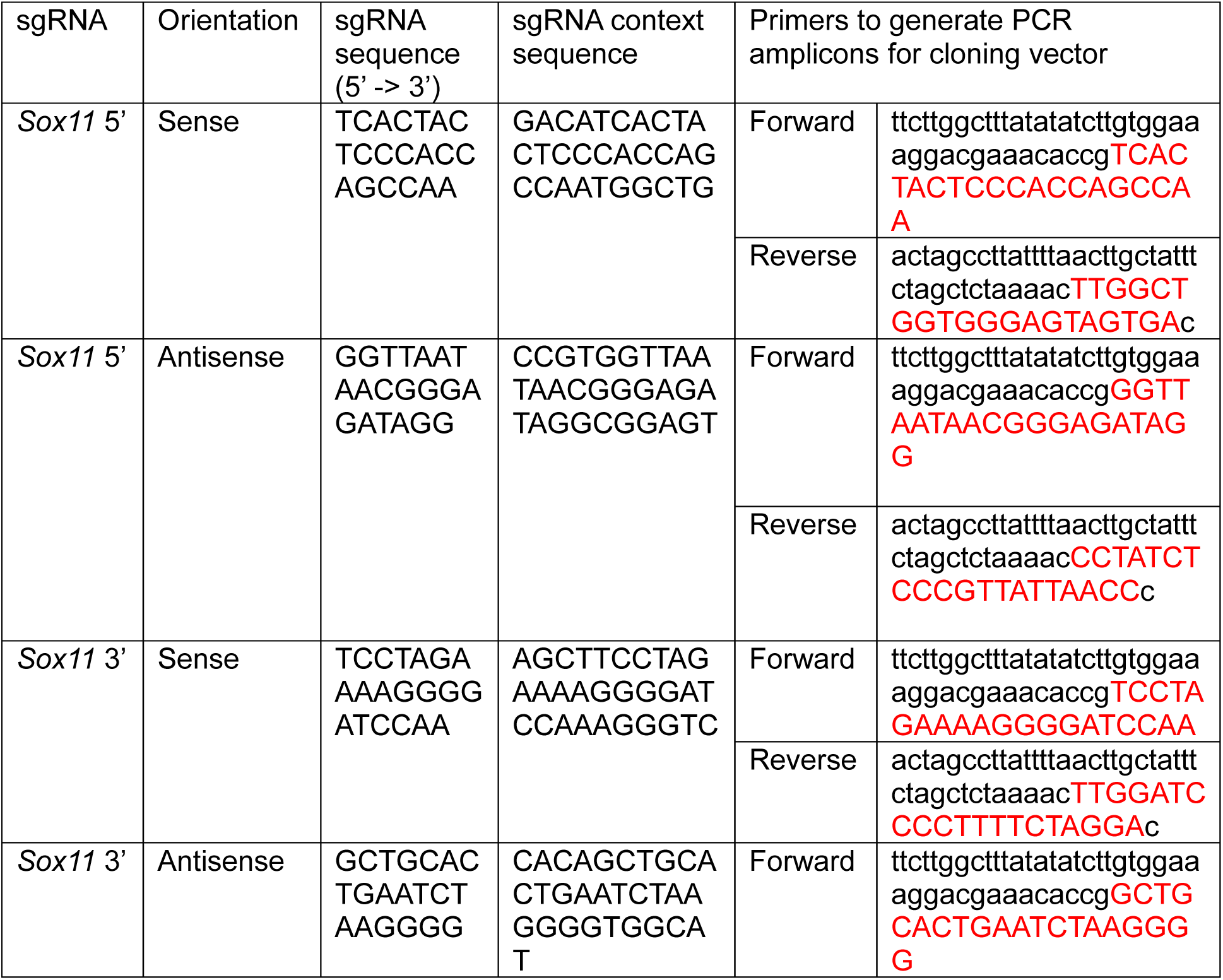
sgRNA sequence cloned into gRNA_Cloning_Vector (Addgene 41824). Red nucleotides represent the sgRNA sequence.

**Table 4:** RNAscope probes.

| **RNAscope Probe** | **Accession Number** | **Probe Region** | **Company** | **Catalog Number** |
| --- | --- | --- | --- | --- |
| Mm-Sox11-O1 | NM_009234.6 | 2956-4380 | Advanced Cell Diagnostics | 538251 |
| Mm-Sox11-O1-sense | NM_009234.6 | 2956-4380 | Advanced Cell Diagnostics | 1212911-C1 |
| Mm-Tubb3-C2 | NM_023279.2 | 2-1636 | Advanced Cell Diagnostics | 423391-C2 |

**Table 5:**
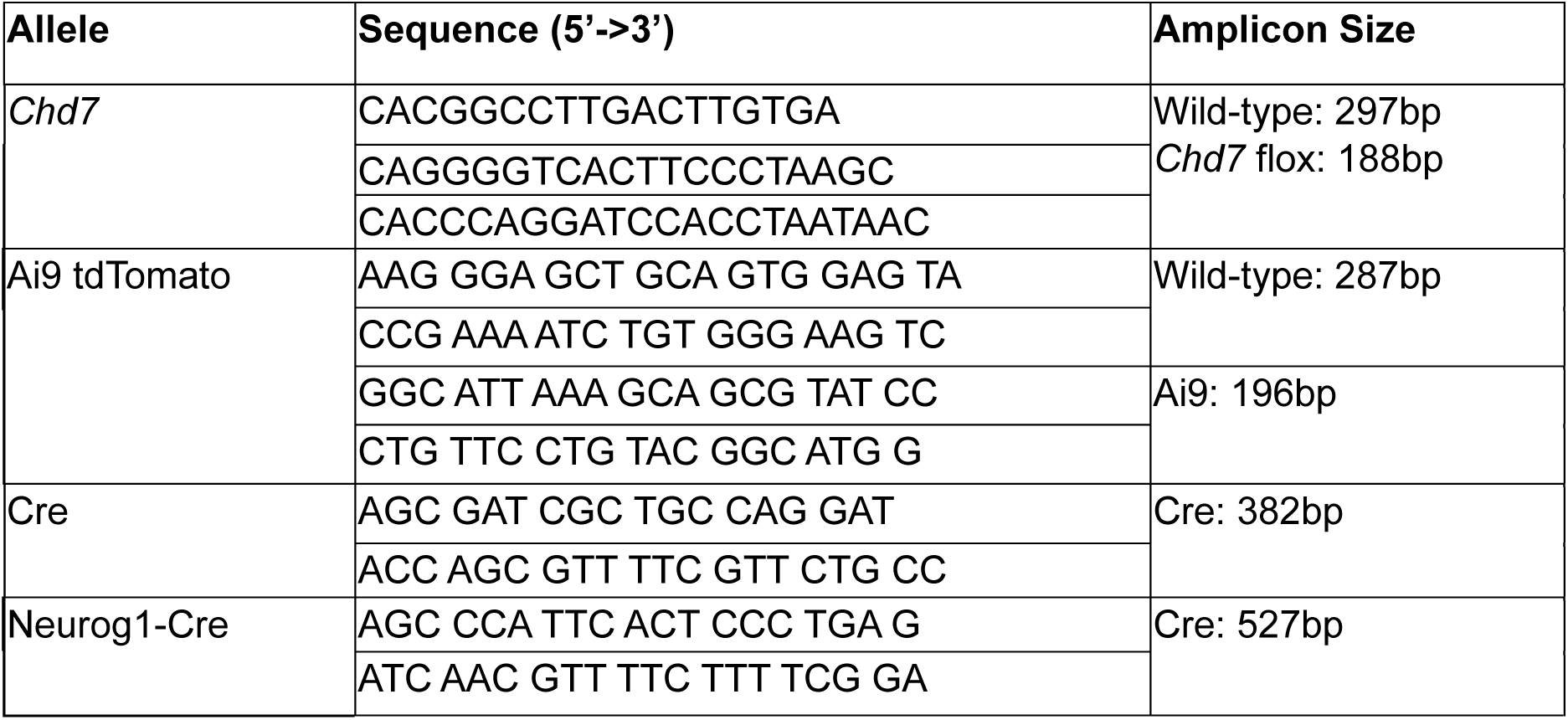
Primers for PCR genotyping.

### CUT&Tag and ChIP-seq library preparation and analysis

Proliferating iMOP or iMOP-derived neurons were gently dissociated using TrypLE Express (Thermofisher) and collected by centrifugation at 1,000 rpm in a clinical centrifuge. Cell numbers were determined using the Moxi Z cell counter (Orflo). 500,000 cells were used for each reaction and processed using the CUT&Tag-IT assay kit (Active Motif, #53160 or #53165). Cells were incubated with either anti-CHD7 (Cell Signaling Technology) or anti-H3K36me3 (Active Motif) antibodies. Antibodies used are listed in Table 1. DNA libraries were prepared from samples, and 150 bp paired-end sequencing was performed on an Illumina NovaSeq S4 Flowcell. CUT&Tag paired-end reads were aligned to the reference genome (mm10) with Bowtie2 (Langmead and Salzberg, 2012). Sam file was converted to bam and then sorted and indexed using Samtools (Li et al., 2009). DeepTools was used to eliminate duplicated reads, and uniquely mapped reads were normalized as Reads Per Kilobase per Million (RPKM) in bigwig files with the bamCoverage (Ramirez et al., 2014). The UCSC bigWigToBedGraph command was used to convert bigwig files into bedgraph (Kent et al., 2010). Sparse Enrichment Analysis for CUT&RUN (SEACR) was used for peak calling(Meers et al., 2019). Bedgraph files from CHD7, EP300, H3K4me3, and CTCF libraries were used as target files, while libraries prepared with only secondary antibodies severed as a threshold for peak calling. Bedtools was used to perform genome arithmetic to identify intersecting regions from CUT&Tag data. ChIP-seq was performed as described by (Kwan et al., 2015). Peak calling was performed using MACS2 (Zhang et al., 2008). Integrative Genomics Viewer (IGV) was used to visualize CUT&Tag (Robinson et al., 2011).

EnrichedHeatmap was used to generate heatmaps and profile plots using signal and region files acquired from CUT&Tag data. Signal files are average bigwig files from two replicates generated using bigwigCompare while region files are the summit regions corresponding to the maximum signals of SEACR peaks. CHD7 average bigwig files were used to obtain the signals, while the summit regions of H3K4me3 (promoter), EP300 (enhancer), and CTCF SEACR peaks were used as regions. Heatmaps of CHD7 signals were centered around +/- 5kb of H3K4me3, EP300, and CTCF summit regions. To visualize CHD7 and EP300 reads at different categories of CHD7-bound enhancers, the summits from common and neuron-specific CHD7-EP300 peaks served as the center. The CHD7 and EP300 average signals were centered around +/- 5kb of the summits. To identify the overlap between CHD7, H3K36me3, and H3K4me3, the intersection regions were identified and acquired from the Gene Expression Omnibus (GEO) database (GSE250033). The rest of the CUT&Tag reads were generated as described in (Kim and Martinez et al., 2024) and were sequenced with the Illumina Hiseq 3000 platform with 150 bp paired-end sequencing.

### RNA-seq library preparation and analysis

Total RNA was extracted from 1-5×10^6^ proliferating or neuronal iMOP cells using 1ml of TRIzol Reagent (Thermo Fisher Scientific #15596026) following the manufacturer’s instructions. Total RNA was treated with DNaseI (NEB #M0303S), and phenol-chloroform extraction was used to remove DNaseI. Following the manufacturer’s instructions, the ribosomal RNA was depleted from 500 ng of DNase-treated total RNA using the NEBNext rRNA depletion Kit (NEB #E6310S). Following the manufacturer’s instructions, the yielded RNA was used for stranded RNA-seq library preparation with the Takara SMARTer Stranded RNA-seq kit (Takara Bio #634838). The average library size, including adaptors, was ∼300bp. Eight uniquely indexed libraries were normalized by concentration and pooled together. Pooled libraries were sequenced using the Illumina Novaseq 6000 S4 platform (2×150bp), resulting in ∼2.5 billion total paired reads.

Sequence reads were mapped to reference mouse genome (mm10) using HISAT2 (Kim et al., 2015). The resulting SAM files were converted, sorted, and indexed to generate sorted BAM files using Samtools (Li et al., 2009). The featureCounts command from Rsubread was used to create the count matrix, and differential expression was calculated using DEseq2 ((Love et al., 2014).To visualize RNA-seq read coverage at forward(+) and reverse(-) strands of the genome separately, we employed Deeptools (3.4.3) (Ramirez et al., 2014) bamCoverage function with sorted BAM files as input.

### RT-qPCR and Strand-specific RT-PCR

Total RNA from proliferative iMOPs and iMOP-derived neurons were isolated using TRIzol reagent (Invitrogen # 15596018) according to the manufacturer’s user guide. For RT-qPCR, 1 µg RNA was used to make cDNA using the qScript cDNA synthesis kit (Quanta Biosciences # 95047-500) according to manufacturer instructions. Relative levels of cDNA were measured by quantitative real-time PCR using the PowerUp SYBR Green Master Mix (Life Technologies #A25742) for 40 cycles of 95°C for 15 s, 60°C for 1 min using the Quantstudio3 real-time PCR machine. Three biological replicates, each with technical triplicates, were used for qPCR. To detect *Chd7* transcripts, the primers 5’ AACCTGTCCTCCACTACAGC 3’ and 5’ TCACTAGCTGAGCGTTCTGT 3’ were used. As a control, *Gapdh* was detected using 5’ AGGTCGGTGTGAACGGATTTG 3’ and 5’ TGTAGACCATGTAGTTGAGGTCA 3’ primers.

For strand-specific RT-PCR, gene-specific primers were used for first-strand cDNA synthesis. Primers complementary to the sense or antisense strands were used for reverse transcription. The oligo dT reaction was used as a positive control, and a no-primer reaction was used as a negative control. The primers and methodology used were based on previously published reports (Ling et al., 2009). 5’ GCACTCGAGTCTGTGAACTAGG 3’ (*Sox11* sense), 5’ GCGTTGTGTGCATAGCAGTC 3’ (*Sox11* antisense), 5’ CCAGGACAATGGCACTGAAT 3’ (*Hmbs* sense), 5’ AAAGTTCCCCAACCTGGAAT 3’ (*Hmbs* antisense) primers were used in individual reactions for first-strand cDNA synthesis with 1 µg of total RNA and ProtoScript II Reverse Transcriptase (New England Biolabs #M0368). Following cDNA synthesis, PCR was performed using the same primer pairs for the respective genes with Taq 5x Master Mix (New England Biolabs #M0285L).

### RNAscope

iMOP-derived neurons used for RNAscope analysis were fixed seven days after initiation of differentiation with 4% formaldehyde in 1X PBS for twenty minutes at room temperature. RNAscope was performed based on the manufacturer’s protocol. iMOP-derived neurons were pretreated with hydrogen peroxide for ten minutes at room temperature, followed by protease III treatment at room temperature for ten minutes.

Probes used for hybridization include Mm-*Sox11*-O1 (Advanced Cell Diagnostics # 538251), Mm-*Sox11*-O1 sense-C1 (#1212911-C1), and Mm-*Tubb3*-C2 (#423391-C2). The Mm-*Sox11*-O1 sense-C1 probe targets the reverse-complement sequence of 2956-4380 bp of ENSMUST00000079063.6. Mm-*Sox11*-O1-sense targets the same region as Mm-*Sox11*-O1 and, therefore, cannot be used together in the same assay. The mRNA within the iMOP-derived neurons was hybridized by incubating with the above probes for 2 hours at 40°C in the HybEZ Oven. Neurons were then incubated with RNAscope amplifier components and tagged with a fluorescent dye. Neurons were incubated at 40°C with AMP1 for 30 minutes, AMP2 for 30 minutes, and AMP3 for 15 minutes. For each channel, cells were treated with channel-specific horseradish peroxidase (HRP-C1, HRP-C2, or HRP-C3) for 15 minutes at 40°C. Next, cells were incubated with either TSA plus fluorescein (Perkin Elmer NEL741001KT), TSA plus Cyanine 3 (Perkin Elmer NEL744001KT), or TSA plus Cyanine 5 (Perkin Elmer NEL745001KT) diluted within TSA buffer for 30 minutes at 40°C. Cells were treated with channel-specific horseradish peroxidase blocker between labeling with each fluorophore for 15 minutes at 40°C. RNAscope reagents used were from the RNAscope Multiplex Fluorescent Detection Kit v2 (Advanced Cell Diagnostics #323110). Following probe hybridization, cells were washed and mounted using ProLong Gold mounting media. For tissue samples, (E)10.5 embryos were collected and fixed by immersion in 4% formaldehyde in 1X PBS overnight at 4°C. Following fixation, tissue samples were washed and cryoprotected with 30% sucrose. 14µm cryosections of E10.5 embryos were collected. Tissue samples were pretreated with RNAscope target retrieval buffer at 98-102°C for five minutes. Following pretreatment, samples hybridized with probes as described above.

### Fluorescence micrograph acquisition and visualization

Samples were mounted on a 1.5-cover glass and acquired on a Zeiss LSM800 point scanning confocal microscope. Antibody-conjugated fluorophores and tdTomato fluorescent protein were excited using one of the four laser lines (405,488, 561, and 633nm) in combination with the appropriate filter sets. 1024×1024 images were acquired with 4X Kalman averaging using either the 20X (0.8 NA) or 40X (1.4 NA) objective. Scale bars were depicted on images.

### *Chd7* conditional knockout mouse

All procedures were based on institutional animal care, used committee research guidelines, and were done at Rutgers University. *Chd7*^flox/flox^ (RRID:IMSR_JAX:030660) mice were crossed with *Neurog1*(*Ngn1*) CreER^T2^ (RRID:IMSR_JAX:008529) and Ai9 R26R tdTomato reporter mice (RRID:IMSR_JAX:007909). Pregnant dams were gavaged with 0.25mg/40g of tamoxifen (Sigma #T5648) and 0.25ug/40g of β-estradiol (Sigma #E8875) daily on E8.5 and 9.5 to induce Cre activity. For staging timed embryos, the morning that vaginal plugs were observed, pregnant female mice were considered carrying E0.5 embryos. Embryos were harvested at the indicated time points.

### Statistical Analysis

All error bars shown in the data are expressed as mean ± standard deviation (sd) of values obtained from independent experiments unless otherwise stated. The numbers (n) of independent experiments were used to perform statistics. Replicate experiments were performed for CUT&Tag, and triplicate experiments were performed for RT-qPCR and RT-PCR. Four independent samples were used for bulk RNA-seq. Statistical analysis was performed with R 4.4.3. The Shapiro-Wilks test was used to determine if the data was normally distributed. Statistical significance was tested in normally distributed data using a two-tailed Student’s t-test. The Wilcoxon rank sum test was used to analyze data that were not normally distributed. For all figures p-values were defined as: * p<0.05, ** p<1×10^-2^, *** p<1×10^-3^, and **** p<1×10^-4^ unless otherwise stated.

## Results

### CHD7 is detectable in proliferating iMOP and iMOP-derived neurons

Proliferating iMOPs and iMOP-derived neurons were used to probe the transcriptome by bulk RNAseq. Proliferating iMOP cells display a transcriptome similar to the primary otic neurosensory cells obtained from E12.5-E13.5 embryonic cochleae (Kwan et al., 2015). The proliferating iMOPs represent the progenitor cell state undergoing self-renewal and do not express cell type-specific markers such as TUBB3 (Fig. 1A). Cells were differentiated into iMOP-derived neurons that express the neuronal marker TUBB3, which is recognized by the Tuj-1 antibody. TUBB3-marked cells have neuronal processes extending out from the soma. Total RNA was harvested from proliferating iMOPs and iMOP-derived neurons and used to generate libraries for bulk RNA-seq. Differential gene expression analysis was performed on bulk RNA-seq, and a volcano plot was used to highlight the differentially expressed genes. In iMOP-derived neurons, *Tubb3* transcripts that code for neuronal β-tubulin were present. *Nhlh1*, a transcription factor involved in developing SGNs, was also identified (Sanders and Kelley, 2022). In contrast, proliferating iMOPs were enriched with proliferation markers such as *Mik67* (Gerdes et al., 1984; Gerdes et al., 1983) and *Top2a* (Nitiss, 1998) and devoid of neuron-specific markers (Fig. 1C). The *Chd7* transcript was detected in both cell types, without significant enrichment in either cell type. To ensure that the transcriptome of iMOP-derived neurons is similar to developing SGNs, we performed a whole transcriptome comparison with E15.5 and P30 SGNs. We used the P8 inner ear glia as a negative control for the comparisons. We obtained the transcriptome data from a published dataset (GSE108620) (Li et al., 2020). The Spearman rank correlation matrix suggested that iMOP-derived neurons were more similar to E15.5 SGNs (σ = 0.72) compared to P30 SGNs (σ = 0.53) (Fig. 1D). In contrast iMOP-derived neurons were inversely correlated with P8 glia (σ = -0.51). The Spearman ranked correlation coefficient ranges from -1 to 1, where 1 is a perfect association and -1 suggests inverse correlation. The results suggest that iMOP-derived neurons have a transcriptome that is similar to early developing SGNs. From this point forward, we refer to proliferating iMOPs as progenitors and iMOP-derived neurons as neurons.

**Fig. 1.**
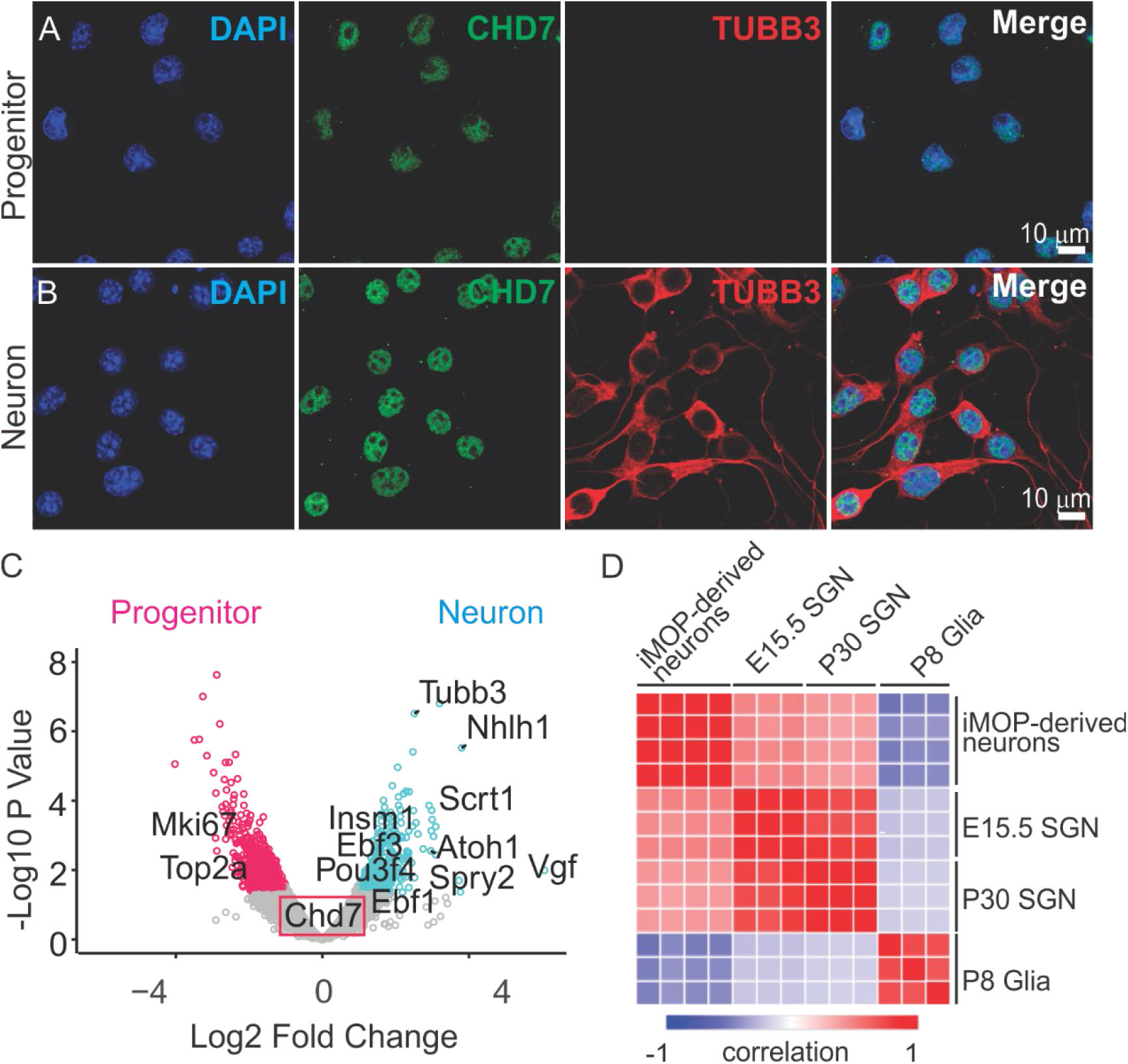
Differential gene enrichment in proliferating iMOP and iMOP-derived neurons. DAPI (blue) and TUBB3 (green) labeling of (A) proliferating immortalized multipotent otic progenitor (iMOP) and (B) iMOP-derived neurons. (C) Differential gene expression of RNAseq from proliferating iMOP (n=4) and iMOP-derived neuron (n=4). Cutoffs were set at p < 0.05 and a log_2_ fold change > 1. Statistically significant enriched genes in iMOP-derived neurons are in cyan, while genes enriched in proliferating iMOPs are in pink. (D) Correlation matrix of transcriptomes from iMOP-derived neurons (n=4), E15.5 (n=3), and P30 spiral ganglion neurons (SGNs) (n=3). P8 glia (n=3) was used as a negative control. SGN and glia data are from (GSE108620). Color represents Spearman rank correlation values, where red represents a high degree of correlation and blue corresponds to anti-correlation.

### Knockdown of *Chd7* disrupts neurogenesis

Although the *Chd7* transcript is detectable, we wanted to determine if CHD7 protein is present. We also wanted to test the importance of CHD7 in neuronal differentiation. We used a knockdown approach to decrease *Chd7* transcripts and dissect the acute effect on neurogenesis. *In vivo* studies of a conditional knockout mouse showed that *Chd7* ablation reduced the number of neuroblasts and altered the expression of proneural genes (Hurd et al., 2010). Control shRNA or a *Chd7* shRNA lentivirus was produced and used to perform the knockdown in progenitors. The transduced cells were cultured in antibiotics to select for infected cells, harvested, and total RNA extracted. RT-qPCR was performed on *Chd7* transcripts from scrambled and *Chd7* shRNA-transduced cells. *Chd7* shRNA transduced cells significantly reduced transcript levels compared to the scrambled shRNA transduced cells (p <0.004, Student’s t-test) (Fig. 2A).

**Fig. 2.**
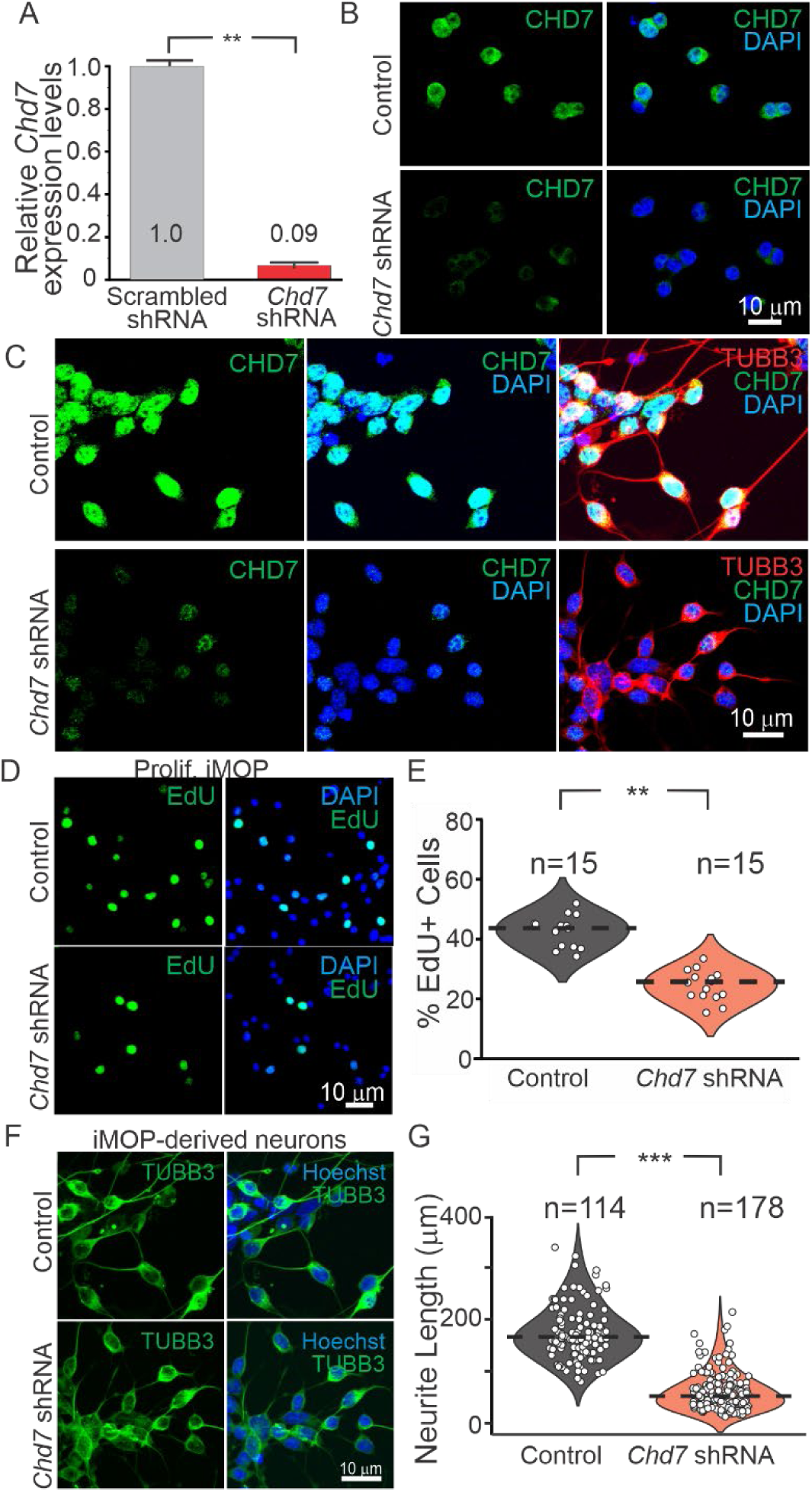
C*h*d7 knockdown affects neurogenesis. (A) *Chd7* transcript levels from RT-qPCR results obtained from scrambled and *Chd7* shRNA stable cell lines (n = 3, independent experiments). (B) Immunofluorescence labeling of CHD7 (green) in DAPI marked nuclei (blue) from control shRNA and *Chd7* shRNA proliferating iMOPs to show knockdown efficacy. (C) CHD7 and TUBB3 immunofluorescence from control shRNA and *Chd7* shRNA iMOP-derived neurons. EdU (green) incorporation in control shRNA and *Chd7* shRNA (D) proliferating iMOPs or (E) iMOP-derived neurons. (F) Quantifying the percent EdU positive cells from control shRNA (n = 3, independent experiments) and *Chd7* shRNA (n=3, independent experiments) proliferating iMOPs. The number of images used for quantification per cell type are listed in the plot. (G) Neurite length measurements from control shRNA (n = 3, independent experiments) and *Chd7* shRNA (n=3, independent experiments). iMOP-derived neurons. Statistics was performed using the number of independent experiments (*** p< 0.001, ** p< 0.01, Student’s t-test).

Decreased CHD7 protein levels were confirmed after knockdown and shown to impair neuronal differentiation. Progenitor and neurons were transduced with lentivirus and subjected to antibiotics selection. Immunostaining against CHD7 and TUBB3 was performed in transduced cells. In progenitors, shRNA control cells showed robust nuclear CHD7 labeling. After transduction with the *Chd7* shRNA virus, the CHD7 protein was barely detectable (Fig. 2B). In control neurons, we observed robust CHD7 expression in the nucleus and TUBB3-marked soma and neurites. In contrast, *Chd7* shRNA-marked neurons had almost undetectable CHD7 levels (Fig. 2C). The data suggested that *Chd7* shRNA dramatically decreases CHD7 protein levels.

Since *Chd7* affected the number of developing neurons *in vivo* (Hurd et al., 2010), we asked whether *Chd7* affects proliferation in our cell culture system. EdU incorporation was performed after *Chd7* knockdown in progenitors. EdU, a nucleotide analog, was added into the culture medium, incorporated into DNA as cells undergo DNA replication and visualized by fluorophore conjugation. Control cells displayed a large percentage of EdU+ cells (42.6 ± 5.24%). In contrast, EdU+ cells decreased in *Chd7* shRNA cells (24.53 ± 5.14%) (Fig. 2D). A statistically significant decrease in the percentage of EdU+ cells was observed after the *Chd7* knockdown (p < 0.003) (Fig. 2E). Since the vast majority of neurons are post-mitotic and express *Cdkn1b*, they did not incorporate EdU as previously shown (Song et al., 2017). Hence, we did not perform EdU incorporation on neurons. To determine the role of CHD7 in neuronal differentiation, control and *Chd7* shRNA transduced neurons were used. The morphological changes in the iMOP-derived neurons were interrogated using TUBB3 immunostaining. TUBB3 marked the soma of the neurons and highlighted the neurites (Fig. 2F). Neurite lengths from control (175.07 ± 52.15µm) and *Chd7* shRNA transduced neurons (60.13 ± 36.5µm) were measured as an indicator of neuronal differentiation. *Chd7* knockdown showed a statistically significant decrease in neurite lengths (p < 0.0004) (Fig. 2G). The results suggest that CHD7 is required for progenitor proliferation and neuronal differentiation during neurogenesis.

### Genome-wide CHD7 enrichment at cis-regulatory elements

Since CHD7 may function in distinct cellular processes in progenitors and neurons, genome-wide enrichment of CHD7 sites was performed in progenitors and neurons. The identified sites provide information on the regions where CHD7 exerts its nucleosome repositioning activity. CUT&Tag was used to determine enrichment sites using several genomic landmarks. For active promoters, H3K4me3 was used to mark active promoters (Benayoun et al., 2014), while EP300 (p300) marked active enhancers (Narita et al., 2021). Finally, CTCF binding sites, which are associated with insulators, were also identified (Huang et al., 2021). Peaks corresponding to significant enrichment of each mark were identified using SEACR. The SEACR-defined peaks corresponded to presumptive promoters, enhancers, and insulator sites. Nucleosome remodeling by CHD7 at the cis-regulatory elements likely imparts an important role in regulating gene expression and chromatin organization in progenitors and neurons. The profile plots and heatmaps show H3K4me3, EP300 and CTCF binding in progenitors. CHD7 displayed similar enrichment levels across these sites (Fig S2A). In neurons, EP300 and CTCF signals were significantly increased. Concomitant to this increase, the CHD7 signal also increased in these regions (Fig. 2SB). Its presence at enhancers agrees with a previous report that showed CHD7 at EP300 and H3K27ac marked enhancers in different cell types (Schnetz et al., 2009). In addition to correlating with enhancer marks, we also identified that CHD7 co-occupied CTCF sites, which could serve as insulator elements. The results suggest that CHD7 may function at cell-type-specific promoters, enhancers and insulators during neuronal differentiation.

### CHD7 enrichment near H3K36me3 sites

Interestingly, H3K4me3 levels were similar in progenitors relative to neurons, but strong H3K4me3 and CHD7 signals were observed outside of the peak region in iMOP-derived neurons. An upset plot of CHD7 enriched sites showed overlap in the promoter, intergenic and distal intergenic regions for both progenitors and neurons (Fig. S3A, B). These regions corresponded to promoters and enhancers. Surprisingly, we observed CHD7 enrichment in the 3’ untranslated regions (3’ UTR) compared to other regions. This overlap suggested that the 3’UTRs may harbor cis-regulatory elements that potentially influence gene expression.

The association of CHD7 at 3’ UTR suggested that it may be enriched at transcriptional end sites (TES) in progenitors and neurons. The H3K36me3 mark was used to determine if CHD7 is enriched at transcriptional end sites. H3K36me3 accumulates at the end of the gene body near the TES. CUT&Tag was performed on H3K36me3, and SEACR-called peaks identified the transcriptional end sites (H3K36me3+) in progenitors and neurons. Peak H3K36me3 signals to define the regions corresponding to the TES. Signals from H3K36me3 and CHD7 were plotted in the regions. H3K4me3, the mark for active promoters, was used as a negative control. In progenitors, the CHD7 signal was observed in H3K36me3 + regions. We did not observe the H3K4me3 promoter mark in most regions since these regions correspond to TES. Surprisingly, we did observe the H3K4me3 signal in a subset of H3K36me3+ regions (Fig 3A). In neurons, a similar enrichment pattern of H3K36me3+ CHD7+ regions and H3K36me3+ H3K4me3+ CHD7+ regions was observed in a different set of sites. The data suggest that CHD7 may function at two distinct sites marked by H3K36me3 alone or regions with H3K4me3 and H3K36me3. The H3K36me3+ mark was recently implicated in DNA damage repair sites and genomic stability maintenance (Sun et al., 2020). In CHD7+ H3K36me3+ regions, CHD7 may facilitate DNA repair and maintain genome stability. However, it is unclear why an active promoter mark may be deposited near the transcriptional end sites.

**Fig. 3.**
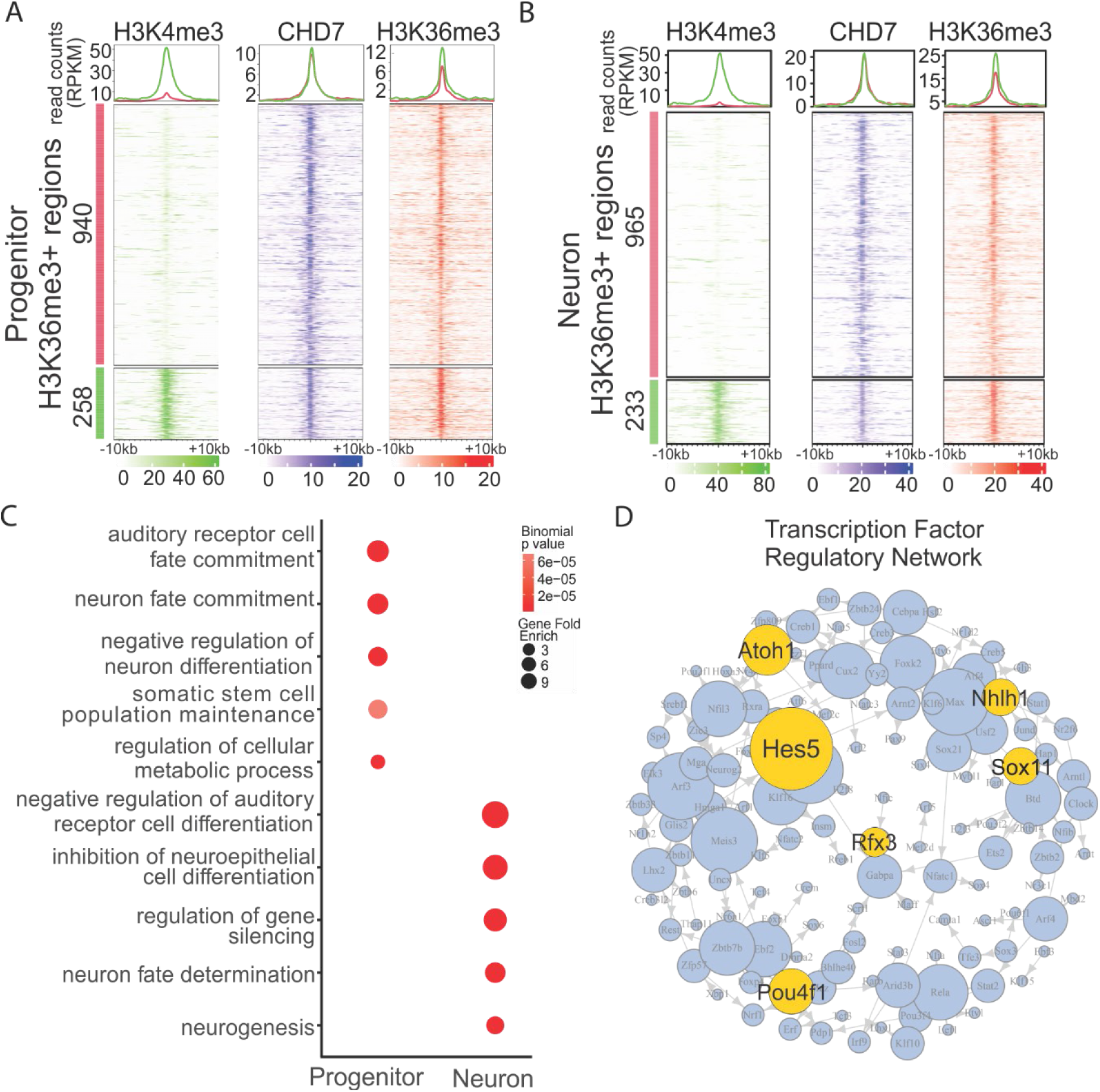
CHD7 enrichment at H3K35me3+ and H3K36me3+ H3K4me3+ sites. H3K4me3 and H3K36me3 were used to define regions corresponding to active promoters and transcription end sites. Hierarchical clustered heatmaps displaying enrichment of CHD7, H3K4me3, and H3K36me3 at (A) proliferating iMOPs and (B) iMOP-derived neurons. (C) Dotplot of GO terms associated with H3K36me3+ CHD7+ H3K4me3+ regions in proliferating iMOPs and iMOP-derived neurons. Dot colors represent statistically significant terms (p < 6 × 10^-5^) with a gene enrichment score of greater than 3. (D) Transcriptional regulatory network analysis using differentially expressed genes from bulk RNA-seq showed predicted gene regulatory networks. Transcription factors are listed within each dot. Predicted interactions between each network is displayed as a connection between dots. Yellow dots highlight terms involved in otic neurogenesis and sensory epithelium development.

### CHD7 is enriched at genes associated with auditory neuron differentiation

A previous report suggested that H3K36me3+ H3K4me3+ sites correspond to the genomic regions that harbor long non-coding RNAs (Guttman et al., 2009). We identified genes near CHD7 binding sites. We performed gene ontology (GO) analysis to gain insight into the biological processes of these genes using GREAT (Gu and Hubschmann, 2023). The H3K36me3+ CDH7+ and H3K36me3+ H3K4me3+ CHD7+ regions from progenitors and neurons were used to identify nearby genes for each set of cells. We used the GO term list to generate a similarity matrix and simplified the gene ontology enrichment results. We focused on GO terms containing the most frequent words identified in the word clouds. In the similarity matrixes for H3K36me3+ CHD7+ sites for progenitors and neurons, a large number of GO terms were associated with metabolic processes, development and differentiation (Fig. S3C, D). For H3K36me3+ CHD7+ H3K4me3+sites, terms for regulatory processes, differentiation and development were observed for the vast majority of GO terms (Fig. S3E, F). GO terms for the H3K36me3+ CHD7+ H3K4me3+ regions were represented as a dot plot for progenitors and neurons. Terms associated with progenitor identity, including auditory receptor cell fate commitment, neuron fate commitment, and somatic stem cell population maintenance, were identified in progenitors. In contrast, terms associated with inhibiting hair cell differentiation (negative regulation of auditory receptor cell differentiation), inhibition of neuroepithelial cell differentiation, along with neuron fate determination and neurogenesis, were identified for neurons (Fig 3C). The results suggest that CHD7 may bind to cell-state dependent H3K36me3+ H3K4me3+ regions to maintain a progenitor fate or promote neuronal differentiation while inhibiting hair cell differentiation.

Gene regulatory networks were used to identify the key regulatory network involved in neuronal differentiation. Gene regulatory networks were inferred from differential gene expression data from RNA-seq (Huynh-Thu et al., 2010). Transcription factors were defined using the JASPAR2024 database. *Nhlh1*, *Pou4f1* and *Sox11* were identified as key regulatory networks in neuronal differentiation and development. *Nhlh1* is a transcription factor involved in early spiral ganglion neuron (SGN) development (Sanders and Kelley, 2022). Similarly, *Pou4f1* has been implicated in cochlear vestibular neuron development (Xu et al., 2024), while *Sox11* is essential for hair cell and SGN development (Gnedeva and Hudspeth, 2015). Other transcription factors associated with hair cell and sensory epithelia development, such as *Atoh1*, Rfx3, and *Hes5,* were also identified (Elkon et al., 2015; Jahan et al., 2013; Su et al., 2015). We interpret that the gene regulatory networks specifying neuronal fate are active while the hair cell networks are repressed as the cells undergo neuronal differentiation. This combined approach of gene ontology and regulatory network analysis helped identify key transcription factors for neuronal differentiation.

We focused on biological processes associated with neurogenesis and neuronal differentiation in iMOP-derived neurons. We surveyed the transcription factors implicated in the associated GO terms from iMOP-derived neurons and refined our candidate to *Sox11*, a key transcription factor required for neurogenesis. *Sox4* and *Sox11* are members of the *SoxC* family of transcription factors detected in the developing sensory organs and spiral ganglion during inner ear development. Deletion of either *Sox* gene stunts inner ear and cochlear neuron development (Gnedeva and Hudspeth, 2015). In other systems, *Sox4* and *Sox11* constitute a gene regulatory network for the progression of neurogenesis (Bergsland et al., 2006).

We looked at the deposition of CHD7, H3K36me3 and H3K4me3 around the *Sox11* gene. Previously, H3K36me3 and H3K4me3 were suggested as histone marks that the identified candidate genes harbor long-noncoding RNAs. Antisense transcripts are long non-coding RNA detected in many species and different cell types (He et al., 2008; Zhang et al., 2006). Antisense transcripts are not evenly distributed across the genome but are found in protein-coding genes with natural antisense transcription (Finocchiaro et al., 2007). Some of the antisense transcripts identified in the 3’UTR preferentially complement the 3’UTR of the target gene (Sun et al., 2005). The complementarity of the antisense transcript with the 3’UTR of the sense transcripts suggested a post-transcriptional role in gene regulation. Although the presence of antisense transcription is well described, it is unclear whether antisense transcripts serve a function or are byproducts of cis-regulatory elements employed during neuronal differentiation.

*Sox4* and *Sox11* harbor antisense transcripts that were temporally regulated during mouse cerebral corticogenesis (Ling et al., 2009). The observation suggests that the antisense transcripts from *Sox4* and *Sox11* correlate to neuronal differentiation. We focused on *Sox11* since the gene regulatory network analysis implicated *Sox11*. *Sox11* is a single exonic gene with a long 3’ UTR. From the CUT&Tag data, CHD7 enrichment was observed around the transcriptional start site of the *Sox11* gene in progenitors and neurons (Site 1). This suggests that CHD7 may affect the chromatin status at the *Sox11* promoter to regulate transcription. In addition to enrichment near the promoter, we saw an increased CHD7 signal at the 3’UTR in iMOP-derived neurons (Site 2 and 3) (Fig. 4A). The data aligns with the finding that CHD7 can bind to the 3’UTR of genes.

**Fig. 4.**
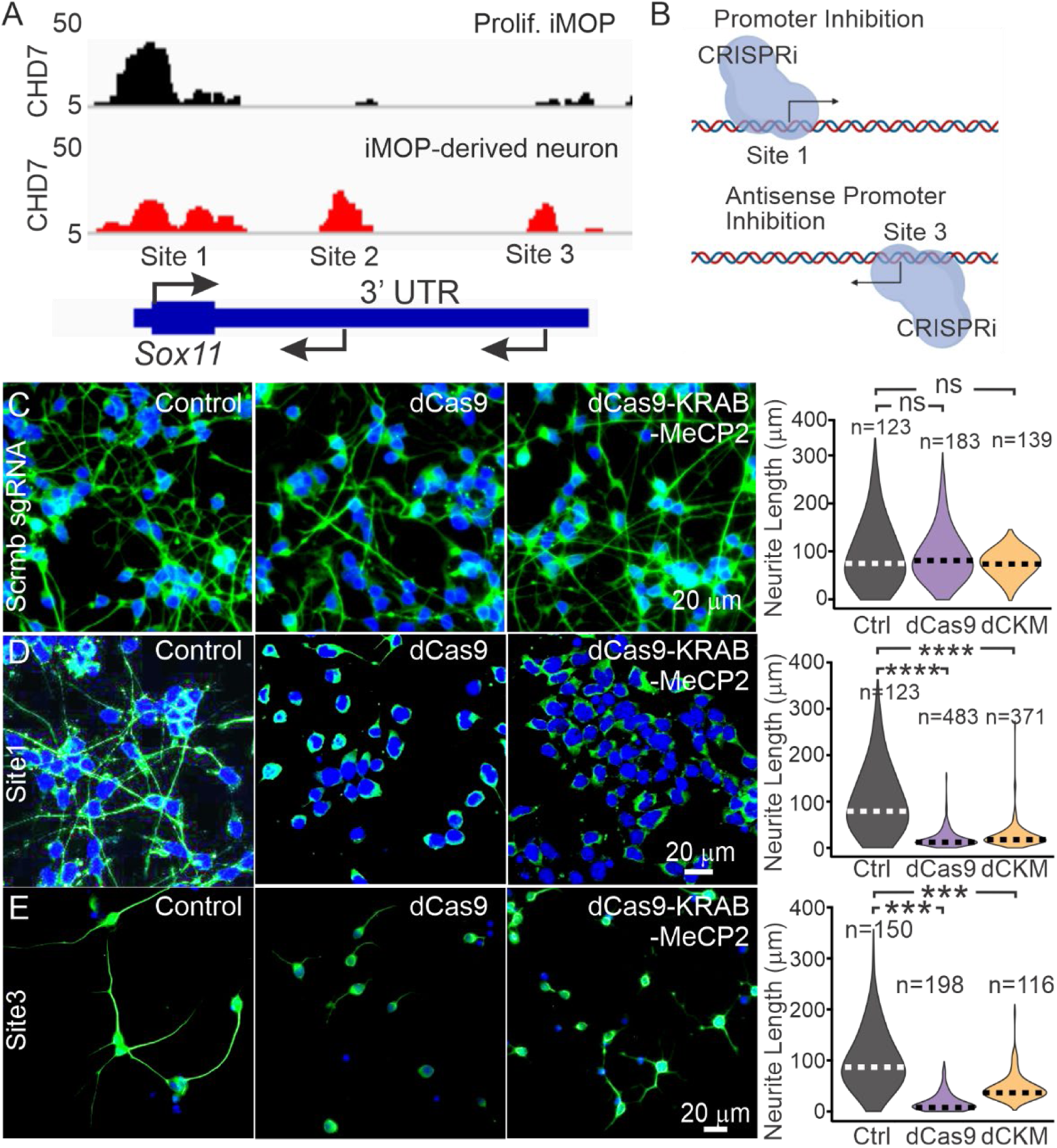
CRISPR inactivation (CRISPRi) of CHD7 sites at the *Sox11* locus. (A) Enrichment of CHD7 around the *Sox11* gene at distinct sites along the *Sox11* locus. (B) CRISPRi inhibition strategy of Site 1, the *Sox11* promoter, and Site 3, a candidate antisense promoter. (C) Constructs containing sgRNAs were transduced into unmodified, dCas9 and dCas9-KRAB-MeCP2 iMOPs. The scrambled sgRNA transduced cell was used as control. sgRNA targeting either Site 1 or Site 3 were used to inactivate the regions. (D) Quantification of neurite lengths in TUBB3 marked cells transduced with scrambled sgRNA or Site 1 sgRNA targeting the *Sox11* promoter. (E) Neurite lengths of a set of iMOP-derived neurons transduced with Site 3 sgRNA. Data were obtained from n=3 independent experiments. The number of cells used for quantification for each stable cell line was listed in the plots. Statistics was performed done using the number of independent experiments (**** p< 0.0001, *** p<0.001; ns,not significant, Student’s t-test).

CHD7 may regulate *Sox11* antisense transcription to affect neuronal differentiation. To delineate the function of CHD7 binding sites in the *Sox11* gene, we sought to block each site using CRISPRi. We used Site 1 as a control to block the promoter, inhibit transcription of *Sox11* and perturb neuronal differentiation. We used Site 3 to represent the presumptive upstream *Sox11* antisense promoter. CRISPRi interfered with Site 1 or Site 3 and was used to determine whether the cis-regulatory element affects neuronal differentiation (Fig. 4B). Lentiviruses harboring dCas9 or dCas9-KRAB-MeCP2 containing the blasticidin resistance cassette were used to transduce iMOP cells to generate stable CRISPRi cell lines. Transduced cells were selected in blasticidin to obtain dCas9 and dCas9-KRAB-MeCP2 stable iMOP cells. The introduction of the sgRNA would direct the dCas9 to prevent protein binding at that region. Similarly, the dCas9-KRAB-MeCP2 complex occludes the site but also employs the KRAB transcriptional repressor domain and a MeCP2 transcriptional repression domain to silence transcription. The combined effect of both repressive domains repressed gene expression compared to dCas9 alone (Yeo et al., 2018). Lentiviruses harboring sgRNA targeting Site 1 or Site 3 that included a neomycin resistance cassette were delivered into either the dCas9 or dCas9-KRAB-MeCP2 stable cell lines, selected in G418 containing medium and differentiated into iMOP-derived neurons for analysis. A scrambled sgRNA transduced cell was used as a control with the dCas9 and dCas9-KRAB-MeCP2 stable iMOP cell line.

We first used the expression of TUBB3 as an early neuronal marker and neurite lengths as a metric for later stages of neuronal differentiation. The percentage of TUBB3+ cells was quantified for each stable iMOP cell line after CRISPRi for Site 1 and Site 3. Interfering with Site 1 with scrambled sgRNA in unmodified cells (93.4 +/- 2.2 %), dCas9 (92.9 ± 8.2% p < 1.00) or dCas9-KRAB-MeCP2 (97.6 ± 2.4%, p <0.51) did not show significant differences in pairwise comparisons with the control (Fig. S3A). In sharp contrast, we observed that interfering with Site 3 decreased the percentage of TUBB3+ cells. Comparing the scrambled sgRNA control (95.9 ± 8.4 %) to either dCas9 (55.9 ± 17.3%, p < 0.049) or dCas9-KRAB-MeCP2 (77.4 ± 22.9%, p < 0.42) showed a significant decrease in TUBB3 expressing cells (Fig. S3B). The data suggest that inhibiting the *Sox11* promoter (Site 1) did not affect the early stages of neuronal differentiation since the percentage of TUBB3-expressing cells remained unchanged. Conversely, interfering with the 3’ UTR (Site 3) affected the early stages of neuronal differentiation because of the significant decrease in the percentage of TUBB3 cells. The data also suggests that the two sites likely have different functions.

Next, we measured neurite lengths in the TUBB3-expressing cells. To ensure that all cell lines have the same differentiation potential, sgRNA was introduced into all stable cell lines and neurite lengths were measured in iMOP-derived neurons. Neurite lengths were normalized relative to the scrambled sgRNA sample. As expected, the median neurite lengths in scrambled sgRNA (80.2 ± 64.7 µm) compared to either dCas9 (86.7 ± 52.5 mm, p = 1.0) or dCas9-KRAB-MeCP2 (100.2 ± 24.6 µm, p = 1.0) did not show significant differences (Fig. 4C). In sharp contrast, many of the neurons after interfering with Site 1 displayed much shorter neurites or no neurites in dCas9 or dCas9-KRAB-MeCP2 cells. Comparing neurite lengths compared to scrambled sgRNA (80.1 ± 64.6 µm,) showed that dCas9 (14.1 ± 18.3 µm, p < 2.88 × 10^-6^) or dCas9-KRAB-MeCP2 (19.9 ± 22.8 µm, p < 8.51 × 10^-6^) cells had significantly reduced neurite lengths (Fig. 4D). Similarly, interfering with Site 3 using dCas9 or dCas9-KRAB-MeCP2, also reduced neurite lengths.

Comparing neurite lengths from sgRNA control (87.1 ± 53.0 µm) to dCas9 (11.6 ± 16.9 µm, p < 2.60 × 10^-4^) and dCas9-KRAB-MeCP2 (38.2 ± 24.7 µm, p < 2.18 × 10^-4^) (Fig. 4E). The results suggest that blocking the *Sox11* promoter (Site 1) or the *Sox11* antisense promoter (Site 3) decreased neurite lengths and inhibited the later stages of neuronal differentiation. The results suggest that inhibiting the 3’ UTR binding site where CHD7 binds is an important cis-regulatory region for early stages of neuronal differentiation distinct from interfering with the *Sox11* promoter.

Previous studies suggested that the *Sox11* antisense transcript may contribute to neuronal development (Ling et al., 2009). To confirm the presence of *Sox11* antisense transcripts, a non-quantitative stranded *Sox11* RT-PCR was done. Total RNA was harvested from proliferating iMOP cells and iMOP-derived neurons and used for strand-specific oligo-mediated reverse transcription followed by PCR amplification. The hydroxymethylbilane synthase gene (*Hmbs*), which does not possess antisense transcripts, served as an internal control (Ling et al., 2009). In addition, a reaction lacking the reverse transcriptase (RT) was used as a negative control. For the sense transcript, we observed a robust PCR amplicon for both *Sox11* and *Hmbs,* as expected. There was no signal for the negative control in progenitors and neurons. The *Sox11* antisense transcripts were observed in both progenitor and neurons, while no detectable antisense transcript was detectable in *Hmbs* or the negative control. Finally, an oligo d(T)_15_ was used as a positive control to amplify the polyadenylated mRNA for *Sox11* and *Hmbs* to ensure RNA integrity (Fig. 5A). The results confirmed the presence of *Sox11* antisense transcripts in progenitors and neurons.

**Fig. 5.**
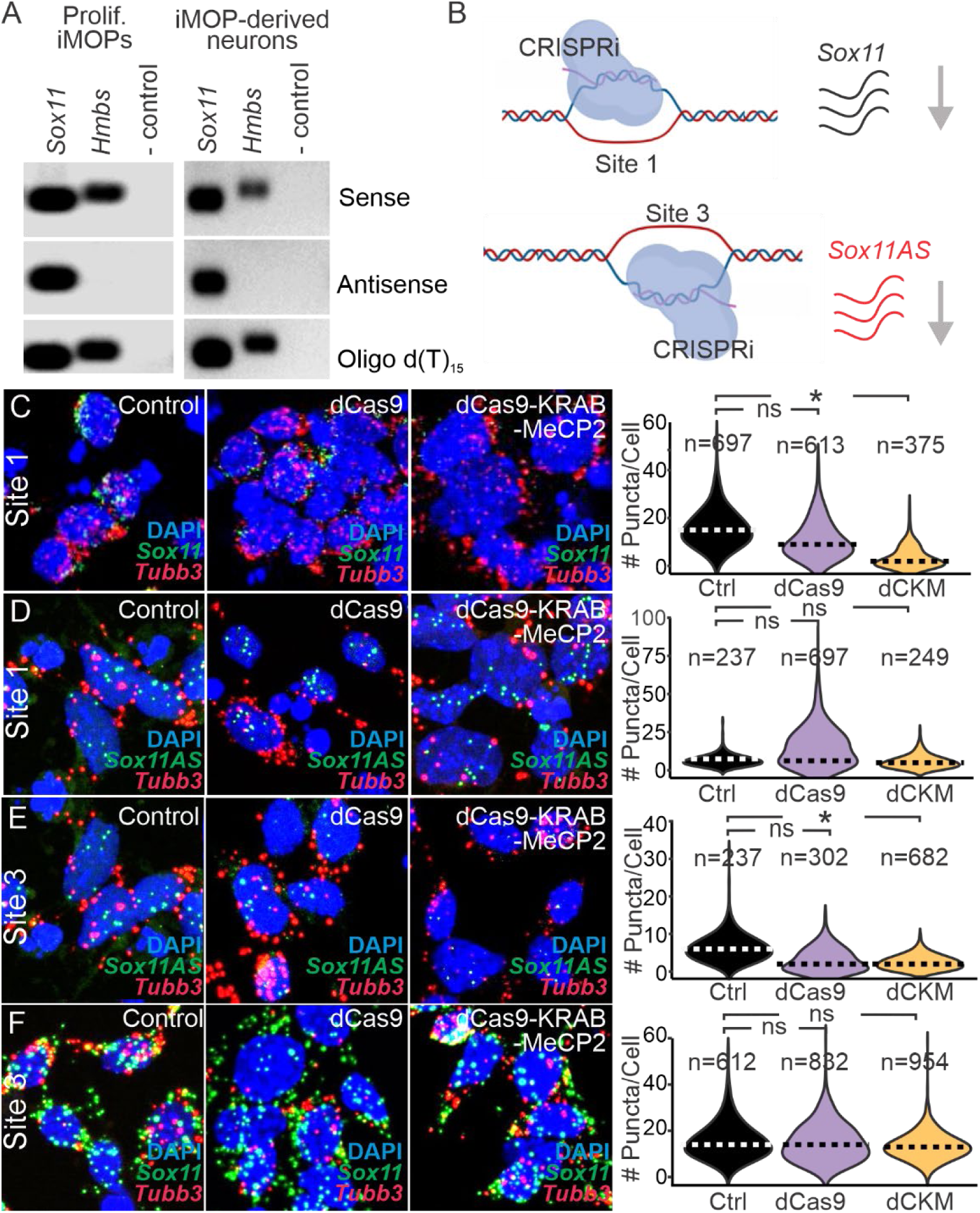
Detection of antisense *Sox11* transcripts after inhibiting candidate antisense promoter. (A) Detection of *Sox11* sense and antisense transcripts in *Sox11*-specific reverse transcriptase reaction. Oligo dT25 reverse transcription served as a control. Hmbs, a gene with no antisense transcript, was used as a control. (B) RNAscope of *Sox11* and *Tubb3* transcripts in DAPI marked nuclei in the cohort of CRISPRi cell lines. *Sox11* antisense (AS) and *Tubb3* transcripts were detected in another group of cells. Quantification of RNA puncta of (C) *Sox11 sense* transcript (control n=3, dCas9 n=3, dCas9-KRAB-MeCP2 n=3, independent experiments) and (D) *Sox11 antisense* transcript in CRISPRi cell cohorts (control n=3, dCas9 n=4, dCas9-KRAB-MeCP2 n=3, independent experiment). RNAscope *Sox11* and *Tubb3* puncta counts after inhibiting Site 3. (E) Quantification of *Sox11* antisense transcripts in *Tubb3* (control n=3, dCas9 n=3, dCas9-KRAB-MeCP2 n=3) independent experiments and (F) *Sox11* sense puncta in *Tubb3* cells (control n=4, dCas9 n=4, dCas9-KRAB-MeCP2 n=4). The number of cells for each stable cell line was listed. Statistics were performed using the number of independent experiments (* p< 0.05; ns, not significant, Student’s t-test)

Although CRISPRi affected neuronal differentiation at CHD7-marked sites, we wanted to clarify whether the *Sox11* antisense promoter (Site 3) could directly or indirectly affect *Sox11* promoter (Site 1) activity. We used the corresponding transcript levels as an indicator for the cis-regulatory element activity (Fig. 5B). Neurons were generated using unmodified (control), dCas9 or dCas9-KRAB-MeCP2 (dCKM) iMOP stable cell line, transduced with lentivirus-expressing sgRNA and selected in antibiotic-containing medium. Single-molecule fluorescence *in situ* hybridization (smFISH) was used to detect and count the relative number of transcripts in individual cells. Quantifying the smFISH puncta allowed us to determine the relative number of transcripts after employing CRISPRi. We designed and used probes to identify *Sox11* and the *Sox11* antisense transcripts. A *Tubb3* probe was used to identify neurons in conjunction with these probes. DAPI-marked nuclei were used to identify cells, and the corresponding puncta were counted in each *Tubb3*-marked cell.

First, we asked whether blocking the *Sox11* promoter (Site 1) decreased *Sox11* transcripts. In control cells, the number of puncta corresponding to *Sox11* sense transcripts (15.85 ± 8.04 puncta/cell) did not significantly decrease in dCas9 cells (10.75 ± 8.01 puncta/cell, p < 0.81), but in dCas9-KRAB-MeCP2 neurons, the *Sox11* puncta (3.38 ± 4.21 puncta/cell, p < 0.043) significantly decreased (Fig. 5C). Puncta quantification confirmed that using dCas9-KRAB-MeCP2 more effectively blocked the *Sox11* promoter (Site 1) to decrease *Sox11* levels. Next, we asked whether the *Sox11* antisense levels were affected after blocking the *Sox11* promoter (Site 1). The number of puncta in control cells (6.33 ± 3.75 puncta/cell) was not significantly different than either dCas9 (5.36 ± 12.79 puncta/cell, p < 0.68) or dCas9-KRAB-MeCP2 cells (5.46 ± 4.21 puncta/cell, p =1.00) (Fig. 5D). These results suggested that inhibiting the *Sox11* promoter (Site 1) does not appreciable affect the *Sox11* antisense promoter (Site 3).

Next, we asked if blocking the *Sox11* antisense promoter (Site 3) decreased *Sox11* antisense levels. Neurons generated from control, dCas9, and dCas9-KRAB-MeCP2 stable cells were transduced with a lentivirus expressing the sgRNA targeting Site 3 and selected in an antibiotic medium. We probed for *Sox11* antisense transcripts in control cells (6.33 ± 3.75 puncta/cell). *Sox11* antisense puncta in dCas9 (2.99 ± 2.98 p < 0.91) did not show a significant decrease compared to the control, but in dCas9-KRAB-MeCP2 cells (2.41 ± 1.85 puncta/cell, p < 0.04),

*Sox11* transcripts were significantly reduced (Fig. 5E). The results suggested that dCas9-KRAB-MeCP2 was more effective in inhibiting the regulatory element. To determine if CRISPRi of Site3 can indirectly affect *Sox11* transcript levels, we compared control to CRISPRi cell lines. In control, *Sox11* puncta was detectable (15.57 ± 7.90 puncta/cell). The puncta numbers remained unchanged when compared to dCas9 (14.80 ± 7.96 puncta/cell, p =1.00) and dCas9-KRAB-MeCP2 cells (13.28 ± 5.72 puncta/cell, p = 1.00) (Fig. 5F). The results suggest that blocking the *Sox 11* antisense promoter (Site 3) decreased *Sox11* antisense levels, and did not affect *Sox11* promoter (Site 1) activity. The results suggest that the *Sox11* and *Sox11* antisense promoters function independently.

### *Chd7* ablation decreases *Sox11* sense and antisense transcripts in otic neurosensory progenitors

We wanted to confirm that CHD7 regulates transcription at the *Sox11* locus *in vivo*. Previous studies using the *Foxg1* Cre driver with two copies of the *Chd7* conditional knockout (cKO) allele resulted in murine embryos with many inner ear phenotypes, including cochlear hypoplasia, absence of semicircular canals, and cristae. To restrict our analysis to developing SGNs, we employed the *Neurog1* CreER^T2^ *Chd7* cKO mice to specifically ask if CHD7 affects transcription of the *Sox11* gene at early stages of developing vestibulocochlear neurons. In the *Neurog1* CreER^T2^ animals, tamoxifen induction of Cre recombinase at embryonic day (E)8.5 marks neurosensory progenitors in the anteroventral quadrant of the developing otic vesicle at E10.5, an early stage of inner ear development. Lineage tracing of the *Neurog1* CreER^T2^ line marked neurosensory progenitors from the delaminating otic vesicle showed that these cells give rise to SGNs (Koundakjian et al., 2007). The *Neurog1* CreER^T2^ transgenic line allows tamoxifen-inducible Cre activity to delete the *Chd7* in the neurosensory progenitors.

We used frozen tissue sections containing otic vesicles from tamoxifen-treated *Neurog1* CreER^T2^ *Chd7* cKO embryos to determine whether CHD7 ablation affects *Sox11* and *Sox11* antisense transcripts during early vestibulocochlear neuron development. The Ai9 tdTomato reporter was introduced into the *Neurog1* CreER^T2^ *Chd7* cKO mice to identify cells with Cre activity. Pregnant dams carrying control and *Chd7* conditional knockout embryos were treated with tamoxifen at E8.5-E9.5, and embryos were harvested at E10.5 when the vestibulocochlear neurons began to delaminate. Tamoxifen induction of Cre in the neurosensory progenitors marked these cells with tdTomato expression. Sections of the inner ear were obtained and used for CHD7 immunostaining. In the fluorescent micrographs, the tdTomato-marked neurosensory cells and delaminating neuroblasts from *Neurog1* CreER^T2^ Ai9 tdTomato control animals identified CHD7 expression, while cell type-specific deletion of *Chd7* using the *Neurog1* CreER^T2^ *Chd7* cKO Ai9 showed a vast majority of the tdTomato marked neurosensory cells and delaminating neuroblasts lacked CHD7 protein (Fig. S6B).

After confirming the lack of CHD7 protein in neurosensory cells, we asked whether *Sox11* and *Sox11* antisense transcript levels were affected in the absence of CHD7 using smFISH. Horizontal cryosections of embryonic tissue containing the developing otic vesicle were identified from E10.5 control and *Chd7* ckO embryos. smFISH was performed on the cryosections containing the otic vesicles using probes that detect *Sox11* and *Sox11* antisense transcripts. tdTomato-expressing cells from the anteroventral region of the otic vesicle from control embryos (Fig. S6C) and *Chd7* cKO otic vesicles (Fig. S6D) were identified. *Sox11* puncta were counted in neurosensory cells from control (Fig. 6A) and *Chd7* cKO (Fig. 6B) embryos. Similarly, for *Sox11* antisense labeling, tdTomato-marked neurosensory cells from control embryos (Fig. S6E) and *Chd7* cKO otic vesicles (Fig. S6F) were identified. *Sox11* antisense puncta were counted from control (Fig. 6D) and *Chd7* cKO (Fig. 6D) neurosensory cells. Quantification showed *Sox11* transcripts (12.24 ± 2.74 puncta/cell) from control significantly decreased in *Chd7* cKO cells (6.18 ± 1.53 puncta/cell, p < 8.24 × 10^-03^) (Fig 6E). *Sox11* antisense transcripts in control (2.78 ± 0.50 puncta/cell) were also decreased in *Chd7* cKO cells (0.75 ± 0.35 puncta/cell, p < 4.41 × 10^-3^) transcripts (Fig. 6F). The results suggest that *Chd7* is essential for regulating both *Sox11* and *Sox11* antisense transcripts *in vivo*.

**Fig. 6.**
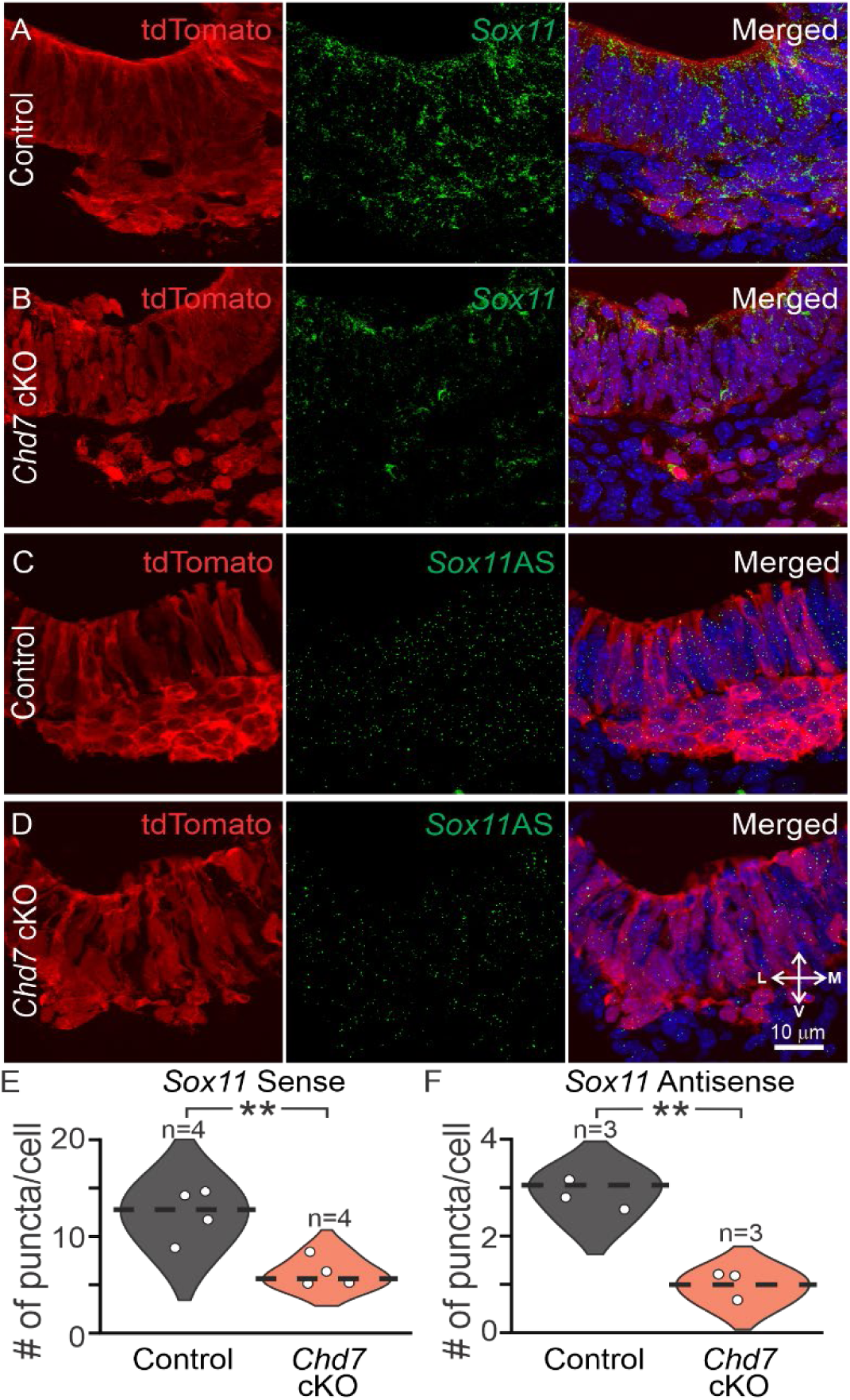
Quantification of *Sox11* and *Sox11* antisense transcripts in otic vesicles of *Chd7* conditional knockout animals. Detection of *Sox11* sense and *Sox11* antisense (AS) transcripts in otic vesicle using RNAscope. (A) Tamoxifen was given to pregnant dams carrying embryos at E8.5-9.5. After tamoxifen treatment, E10.5 *Neurog1* CreER^T2^ Ai9 tdTomato (Control) and (B) *Neurog1* CreER^T2^ *Chd7*^floxed/floxed^ Ai9 tdTomato (*Chd7* cKO) were harvested. Sections of the otic vesicle were used for RNAScope to quantify *Sox11* transcripts from (A) Control and (B) *Chd7* cKO cells. Similarly, RNAscope was performed to identify *Sox11* antisense transcripts in (C) Control and (D) *Chd7* cKO tdTomato marked cells. tdTomato-marked neurosensory cells from the otic vesicle were used for quantification. (D) *Sox11* transcripts displayed a marked decrease comparing Controls (n=4 animals, 8 otic vesicles) to *Chd7* cKO cells (n=4 animals, 7 otic vesicles, p <8.24 × 10^-3^). (E) Similarly, comparison of *Sox11* antisense transcripts from Controls (n=3 animals, 6 otic vesicles) to *Chd7* cKO (n=3 animals, 5 otic vesicles, p < 4.41 × 10^-3^) cells showed a significant reduction. Statistics was performed by the number of animals.

### CHD7 co-occupied CTCF sites at TAD boundaries

Since the antisense promoter did not affect *Sox11* transcript levels, we ruled out its function as a cis-regulatory element that affects *Sox11* transcription. Our genome-wide co-occupancy data showed that CHD7 can co-occupy CTCF sites. CTCF binding can mark putative cell type-specific insulators that prevent the spread of chromatin state between topologically associating domains (TADs). Insulator elements, marked by CTCF, are often located at the boundaries of TADs, the fundamental units of three-dimensional genome organization. CTCF plays a crucial role in defining TAD boundaries, which are primarily conserved across cell types and species (Dixon et al., 2012; Rao et al., 2014). We utilized Hi-C data from cortical neurons to define neuronal TADs (Bonev et al., 2017). We showed that *Sox11* resides at a TAD boundary (Fig. 7A). We compared our CTCF+CHD7+ peaks to TAD boundaries. We observed that the CTCF+ CHD7+ site corresponding to the *Sox11* antisense promoter resides within the boundary element. Genomic tracks of CTCF and CHD7 enrichment centered around the *Sox11* gene confirmed the presence of the proteins at the TAD boundary (Fig. 7B). We also observed that members of the *SoxC* family of transcription factors involved in neurogenesis (Wegner, 2011) also harbored potential insulators. *Sox4* (Fig. S5A) and *Sox12* (Fig. S5B) resided at TAD boundaries and displayed CHD7 and CTCF marks around their loci. We propose that CHD7 functions at insulators between conserved TADs, and disruption of these sites, such as the *Sox11* 3’UTR, affected neuronal differentiation.

**Fig. 7.**
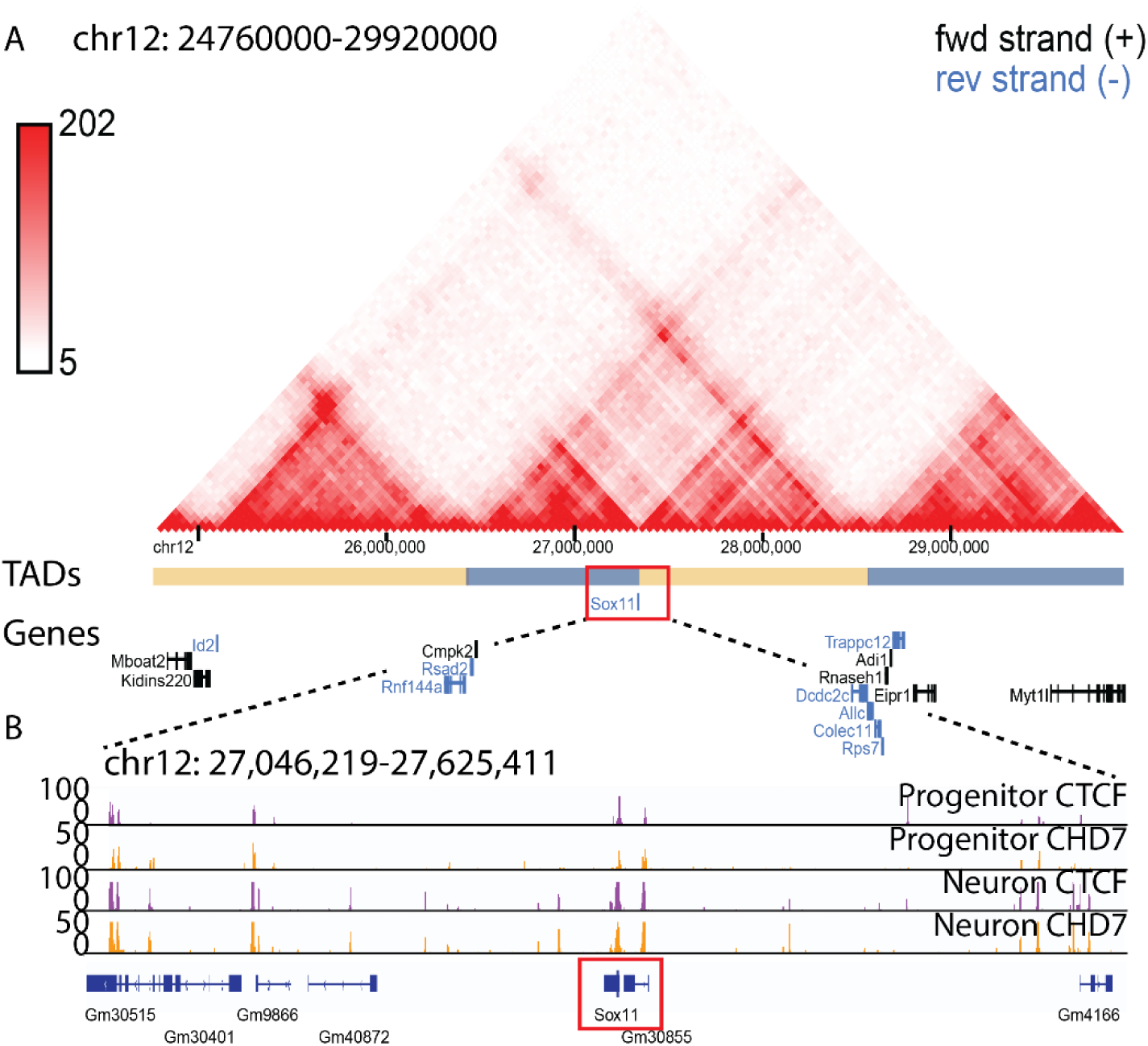
Topologically associating domains around the *Sox11* locus. (A) Contact maps for topologically associating domains (TADs) centered at the *Sox11* gene. from cortical neurons (Bonev et al., 2017) were visualized using the 3D Genome Browser (Wang et al., 2018). For simplicity, many of the non-coding genes were not shown. Predicted TAD regions were indicated by blue and yellow bars. *Sox11* resides at a boundary region between a pair of TADs. (B) A zoomed in view of CTCF and CHD7 peaks around *Sox11.* Genomic tracks displayed CTCF+ CHD7+ peaks in both progenitors and neurons. Genomic coordinates for TADs and genomic tracks as labeled.

## Discussion

### Neurogenesis requires *Chd7* in differentiating iMOPs

We showed that iMOPs were transcriptionally similar to SGNs and could be used to study aspects of SGN development. iMOPs were derived from SOX2 expressing progenitors from E12.5-E13.5 embryonic cochlea (Kwan et al., 2015). The iMOP-derived neurons displayed a transcriptome that was similar to an early developmental time for SGN development. The cells provide a platform for understanding how SGN differentiation occurs. Differential gene expression analysis suggested that key transcription factors such as *Nhlh1* and *Neurod1* were increased in iMOP-derived neurons. *Neurod1* is a well-described transcription factor that is necessary for the survival and development of SGNs. NeuroD1 is essential for SGN development (Evsen et al., 2013; Liu et al., 2000), and conditional deletion of *Neurod1* resulted in SGN degeneration and innervation deficits in the peripheral and central axons (Jahan et al., 2010). *Nhlh1* is expressed in unspecialized SGNs and is rapidly downregulated in single-cell RNAseq data after differentiating into specific subtypes of trajectory inference (Petitpre et al., 2022). The results suggest that iMOPs may express *Nhlh1* and employ the transcription factor for the initial steps that guide unspecialized neurons into the different SGNs subtypes. The iMOP cells can inform future work on SGN development and be used to generate SGN subtypes.

*Chd7* is required for neurogenesis in iMOPs based on the reduced percentage of EdU-marked proliferating progenitors and decreased TUBB3-expressing neurons after *Chd7* knockdown. Even when cells express the neuronal marker TUBB3, the cells display shortened neurites. These results align with observations from *in vivo* studies*. Chd7* conditional knockouts displayed cochlear hypoplasia and the absence of semicircular canals and cristae. Furthermore, there were notable changes in the vestibulocochlear ganglion neurons. *Chd7* deficiency reduced the number of developing neuroblasts, decreased the number of NEUROD1-expressing neuroblasts, and altered proneural gene expression (Hurd et al., 2010). Our findings using iMOPs support CHD7’s function in cellular processes involved in otic neurogenesis.

### Regulation of Transcriptional Regulatory Networks

Prediction of the transcriptional regulatory network highlighted several key transcription factors, such as *Sox11*, *Nhlh1* and *Pou4f1*, in the neuronal differentiation of iMOP-derived neurons. *Sox11* is expressed in SGN and other inner ear cell types. Ablating *Sox11* and the remaining *SoxC* family of transcription factors results in inner ear organ hypoplasia (Gnedeva and Hudspeth, 2015) and may affect SGN development. *Nhlh1* has been identified in multiple single-cell RNA sequencing datasets and is proposed to facilitate the specialization of unspecialized neurons in the cochlea (Petitpre et al., 2022). *Pou4f1* (POU Class 4 Homeobox 1) is a transcription factor that is broadly expressed at birth and essential for SGN development (Xu et al., 2024). The presence of the aforementioned transcriptional regulatory network fits with unspecialized otic neuronal identity before they differentiate into four distinct SGN subtypes. In mice, the type Ia, Ib, Ic, and type II neurons constitute the neuronal subtypes with unique properties contributing to sound detection. Type I SGNs primarily innervate inner hair cells and transmit most auditory information to the brain (Grandi et al., 2020; Petitpre et al., 2022; Shrestha et al., 2018; Sun et al., 2022), while type II unmyelinated SGNs are unmyelinated neurons detect noise damage (Flores et al., 2015; Liu et al., 2015). The analysis suggested that many gene regulatory networks in unspecialized developing SGNs are present in iMOP-derived neurons. Additionally, transcription factors involved in cochlear sensory epithelium and hair cell development, such as *Atoh1*, *Hes5*, and *Rfx3*, play critical roles. *Atoh1* is essential for hair cell development, with its deletion resulting in the absence of hair cells (Bermingham et al., 1999). *Hes5* suppresses hair cell identity in supporting cells, establishing the mosaic of hair cells and supporting cells in the sensory epithelium (Zine et al., 2001). *Rfx3* is crucial for inner ear hair cell development and function, with its loss leading to profound hearing loss (Elkon et al., 2015). Components of these transcriptional regulatory units are present but likely repressed to inhibit hair cells and supporting cell differentiation to facilitate neuronal differentiation in otic progenitors. Our data suggested that iMOPs retain many transcriptional regulatory networks for hair cells, supporting cell and otic neuron development.

### Identifying CHD7-enriched cis-regulatory regions involved in neurogenesis

Harvesting SGNs for genome-wide binding studies is difficult due to the inaccessibility of the inner ear and the small number of harvested neurons. Genome-wide binding studies using iMOPs suggest that CHD7 was enriched at different cis-regulatory elements. We used iMOPs to identify novel regulatory elements that require CHD7 function. We identified CHD7 binding at H3K36me3 sites where it could function in DNA repair. We also identified novel regions marked by H3K4me3 and H3K36me3. We interpret that the regions harboring H3K4me3 and H3K36me3 did not simultaneously occupy the sites but represented different cellular states. The H3K36me3+ H3K4me3+ CHD7+ regions corresponded to regulatory elements that transcribe non-coding RNA.

### CHD7 binds to H3K36me3+ H3K4me3+ sites that transcribe non-coding RNA

Gene ontology analysis using the H3K36me3+ H3K4me3+ CHD7+ regions identified key transcriptional regulatory networks, such as *Sox11*, that affect neurogenesis. We wanted to validate these targets functionally and used the *Sox11* locus as an example. We identified that CHD7 binds to *Sox11* promoter and the 3’UTR of *Sox11*. The data suggested two distinct mechanisms of CHD7 function at the *Sox11* locus. *C*HD*7* regulates *Sox11* transcription by modulating the epigenetic status of the promoter during neuronal differentiation. However, it is unknown how CHD7 functions at the 3’UTR of *Sox11*. Since there are overlapping sequences between the two transcripts, the transcripts may form a double-stranded RNA. The double-stranded RNA may elicit a DICER-mediated degradation that decreases *Sox11* transcript levels.

However, our data did not support this idea. Reducing the *Sox11* antisense transcript levels by blocking its corresponding promoter region did not appreciably affect *Sox11* levels. The results do not rule out that the antisense transcript may still affect SOX11 protein levels. Instead, we propose that the *Sox11* antisense transcript is a byproduct of a cis-regulatory element in the *Sox11* 3’ UTR. Our *in vivo* studies using the *Chd7* cKO animals confirm that CHD7 affects both *Sox11* and *Sox11* antisense levels in otic neurosensory progenitors. We propose that CHD7 contributes to neuronal differentiation in two ways: first, by directly regulating the transcription of *Sox11*, and second, by regulating the function of a regulatory site in the 3’ UTR of *Sox11*.

### CHD7 co-occupied CTCF sites and resided at TAD boundaries

CTCF is demonstrated to generate chromatin looks and organize the loops into topologically associated domains (TADs). CTCF marks 3’UTR of Sox11 and resides at a TAD boundary. Inhibiting a singular site at the 3’UTR of *Sox11* reduced the percentage of TUBB3-expressing neurons and shortened neurites. The data suggests that the singular site corresponds to an insulator element that significantly contributes to neuronal differentiation instead of functioning as an antisense promoter to generate a *Sox11* antisense transcript. We propose that the CHD7+ CTCF+ site produces non-coding RNA as a byproduct, which may serve as a signature of a functional regulatory element. We suggest that CRISPRi displaces CHD7 from the site during neuronal differentiation and prevents CHD7’s nucleosome activity from establishing a functional insulator that separates the TADs. Promiscuous enhancer and promoter interaction between TADs may occur without a boundary and affect gene expression. Altered gene expression would ultimately disrupt cellular processes such as neuronal differentiation.

Our study identified novel regulatory regions bound by CHD7 that regulate neurogenesis. From genome-wide CUT&Tag data, we identified CHD7 binding to promoters, enhancers and insulators. Validating one of these sites marked by CHD7 and CTCF suggests that the region can harbor H3K36me3 and H3K4me3 and produce non-coding RNA, which we dub insulator RNA. The production of non-coding RNA likely corresponds to RNA polymerases in the macromolecular complexes recruited to the region. Interfering with a single region located at the 3’ UTR of *Sox11*, dramatically interfered with neurogenesis. We propose that the site be an insulator element separating TADs to organize the 3D chromatin structure. Disruption of the cis-regulatory element would perturb chromatin organization and affect neurogenesis. CHD7 may function at similar sites scattered throughout the genome to organize chromatin structure. A reduction in CHD7 activity, as proposed for CHARGE syndrome, preferentially affects conserved TADs required for developing inner ear cell types. The increased probability of promiscuous contact inter-TAD regions may perturb the developmental process to varying degrees and explain the variable phenotype observed for patients with CHARGE.

## Author contributions

EM, AJ, JN, JQ, JK and KYK performed the experiments and interpreted and analyzed the data. JQ, JN, and KYK performed bioinformatics analysis. EM and KYK wrote the manuscript.

## Accession numbers

Sequence files were deposited in GEO. RNA-seq fastq files, raw counts and RPKM counts are available in GSE293437. ChIP-seq fastq files and processed bigwig obtained from MACS2 can be obtained from GSE293438. CUT&Tag fastq files, processed bigwig and SEACR peak files are available in GSE293439.

## Acknowledgments

R01 DC018404 supports KYK. KYK acknowledges previous support from the Duncan and Nancy MacMillan Faculty Development Chair Endowment Fund, Busch Biomedical Research Grant, Rutgers Faculty Development Grant, CHARGE Foundation Grant, and R01 DC015000.

## Supplemental Figures

**Fig S1.**
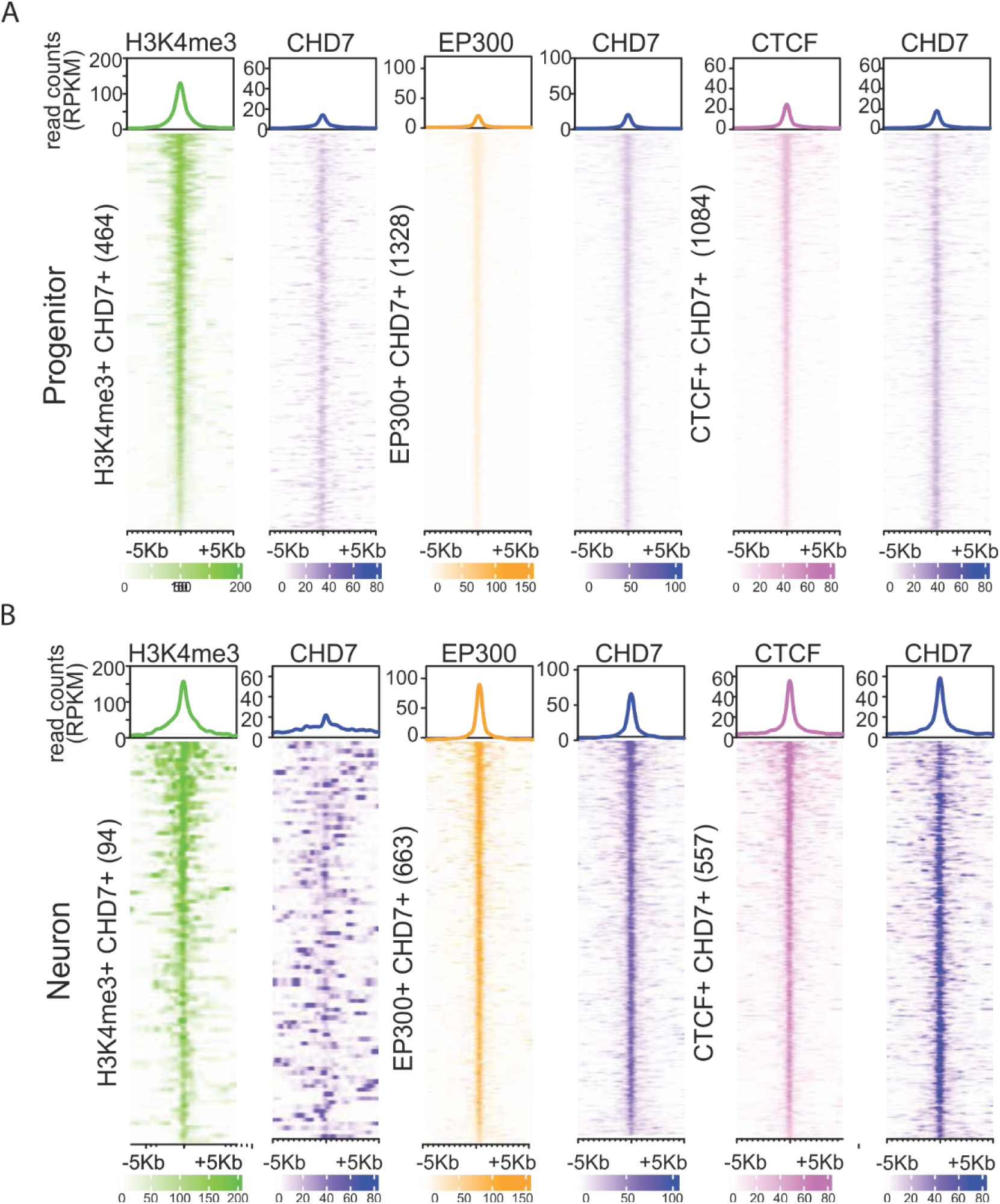
CHD7 enrichment at regulatory elements. SEACR peaks from CUT&Tag of H3K4me3, EP300, and CTCF were used to define distinct regulatory sequences. CHD7 enrichment with each of the regulatory marks was used to determine the regions of interest. The number of CHD7-associated regulatory sites H3K4me3+ CHD7+, EP300+ CHD7+, and CTCF+ CHD7+ were stated. Heatmap and profile plots were centered around these regions in a +/- 5kb window. Signals from CHD7 and regulatory marks were plotted from (A) proliferating iMOPs and (B) iMOP-derived neurons.

**Fig. S2.**
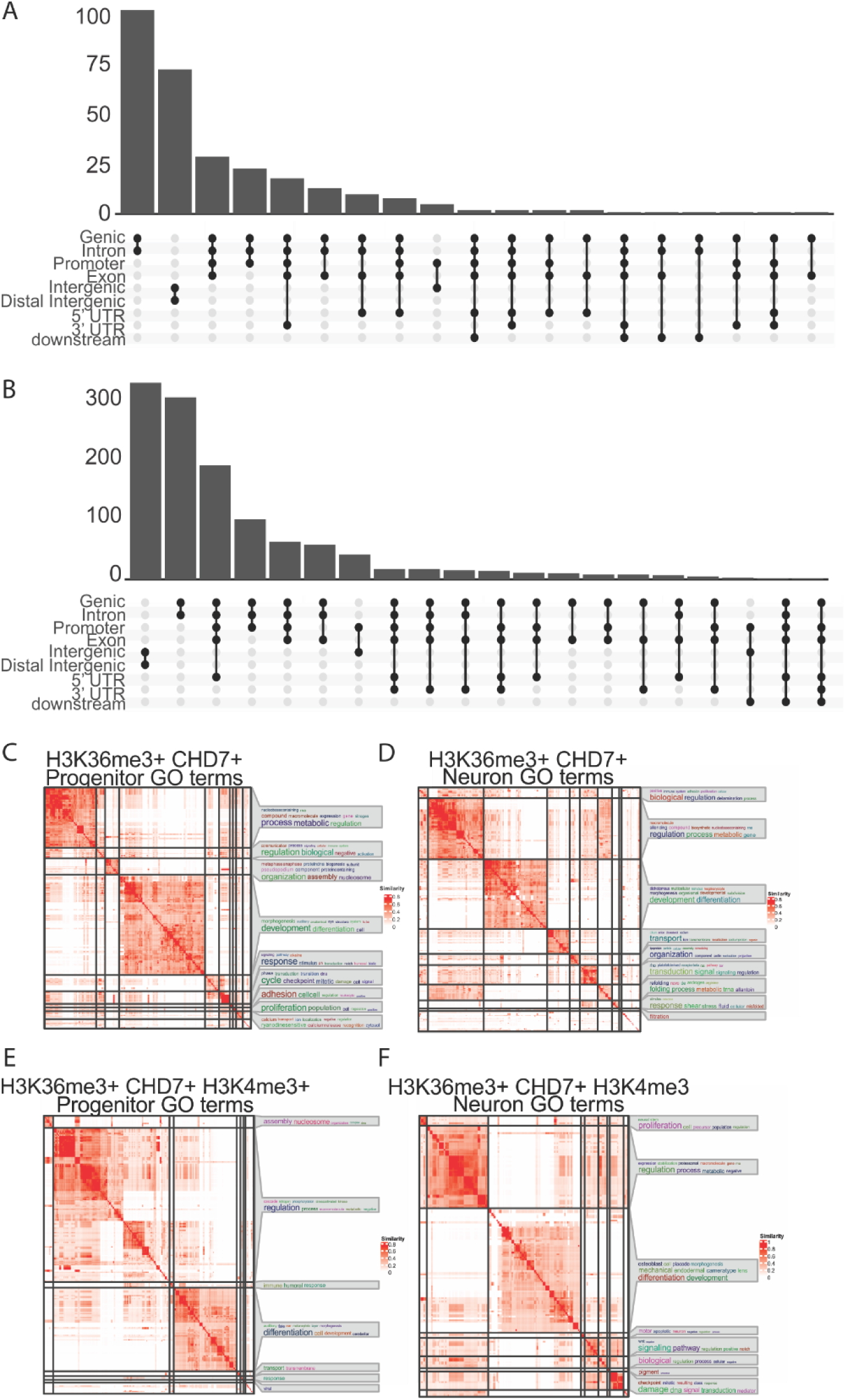
Gene Ontology analysis on H3K36me3+ CHD7+ and H3K36me3+ H3K4me3+ CHD7+ sites. Upset plots of (A) H3K36me3+ CHD7+ and (B) H3K36me3+ H3K4me3+ CHD7+ regions showing the overlap of 3’UTR binding to other regions. GO analysis using regions obtained from CUT&Tag. GO terms were clustered into similar groups and displayed as a matrix using the simplifyEnrichment package. Word clouds on the right of the matrixes display the key words that are over-represented in each category. (A) H3K36me3+ CHD7+ sites in progenitors and neurons displayed terms associated with metabolism, proliferation, development and differentiation (B) H3K36me3+ H3K4me3+ CHD7+ sites in progenitors and neurons were also associated with regulatory processes, development and differentiation.

**Fig. S3.**
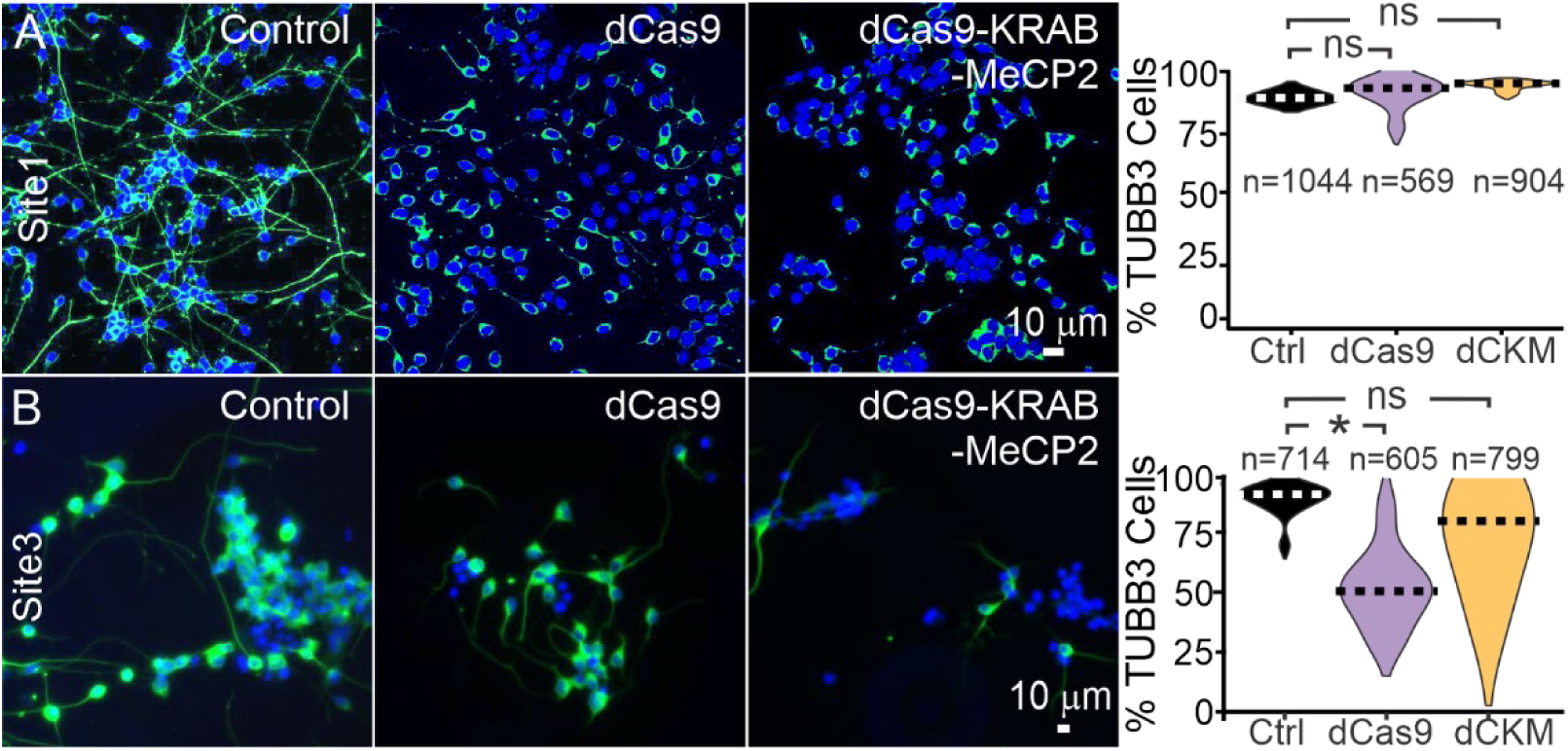
Percentage of TUBB3 cells after sgRNA. The Site 1 sgRNA was used to transduced iMOP, dCas9, and dCas9-KRAB-MeCP2 stable cell lines were immunostained with TUBB3 antibodies. The percentage of TUBB3 marked cells were quantified. Violin plots showing percentage of TUBB3 expressing cells from each cell line. The number of cells quantified for each cell line were denoted. (A) For CRISPRi of Site 1, control (n=4), dCas9 (n=4) and dCas9-KRAB-MeCP2 (n=4 independent experiments were performed. No significant (ns) changes in TUBB3 percentages were observed when blocking Site 1. (B) For CRISPRi of Site 3, control (n=3), dCas9 (n=3) and dCas9-KRAB-MeCP2 (n=3) independent experiments were performed. A significant difference in the percentage of TUBB3 cells was observed after blocking Site 3 in dCas9 (p < 0.049), but the dCas9-KRAB-MeCP2 cells did not show a significant change (p < 0.42). The number of independent experiments was used for Student’s t-test.

**Fig. S4.**
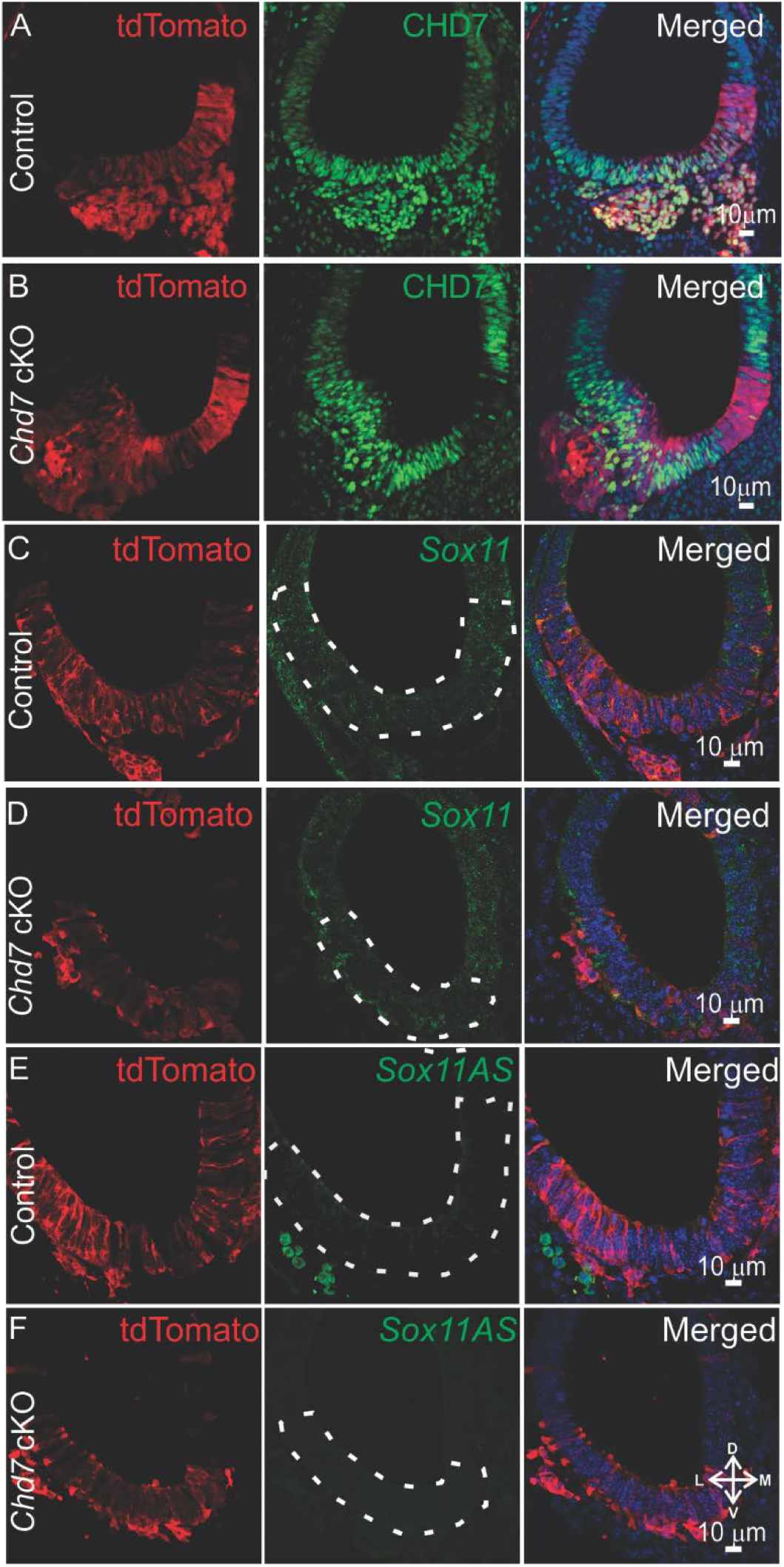
Ablation of *Chd7* from neurosensory domain of the otic vesicle. *Neurog1*-CreER^T2^ Ai9 tdTomato (control) and *Neurog1*-CreER^T2^ *Chd7*^flox/flox^ Ai9 tdTomato (*Chd7* cKO) embryos were harvested at E10.5. *Neurog1*-Cre is expressed in neurosensory cells and marks cells with the red fluorescent protein, tdTomato. Frozen sections containing the otic vesicle from (A) Control tdTomato expressing cells from the neurosensory region of the otic vesicle express CHD7. (B) In *Chd7* cKO embryos, CHD7 protein is no longer present in vast majority of tdTomato marked neurosensory cells. RNAScope was performed to detect *Sox11* transcripts in from (C) Control and (D) *Chd7* cKO cells. Detection of *Sox11* antisense transcripts from (C) Control and (D) *Chd7* cKO cells. tdTomato marked neurosensory cells are outlined.

**Fig. S5.**
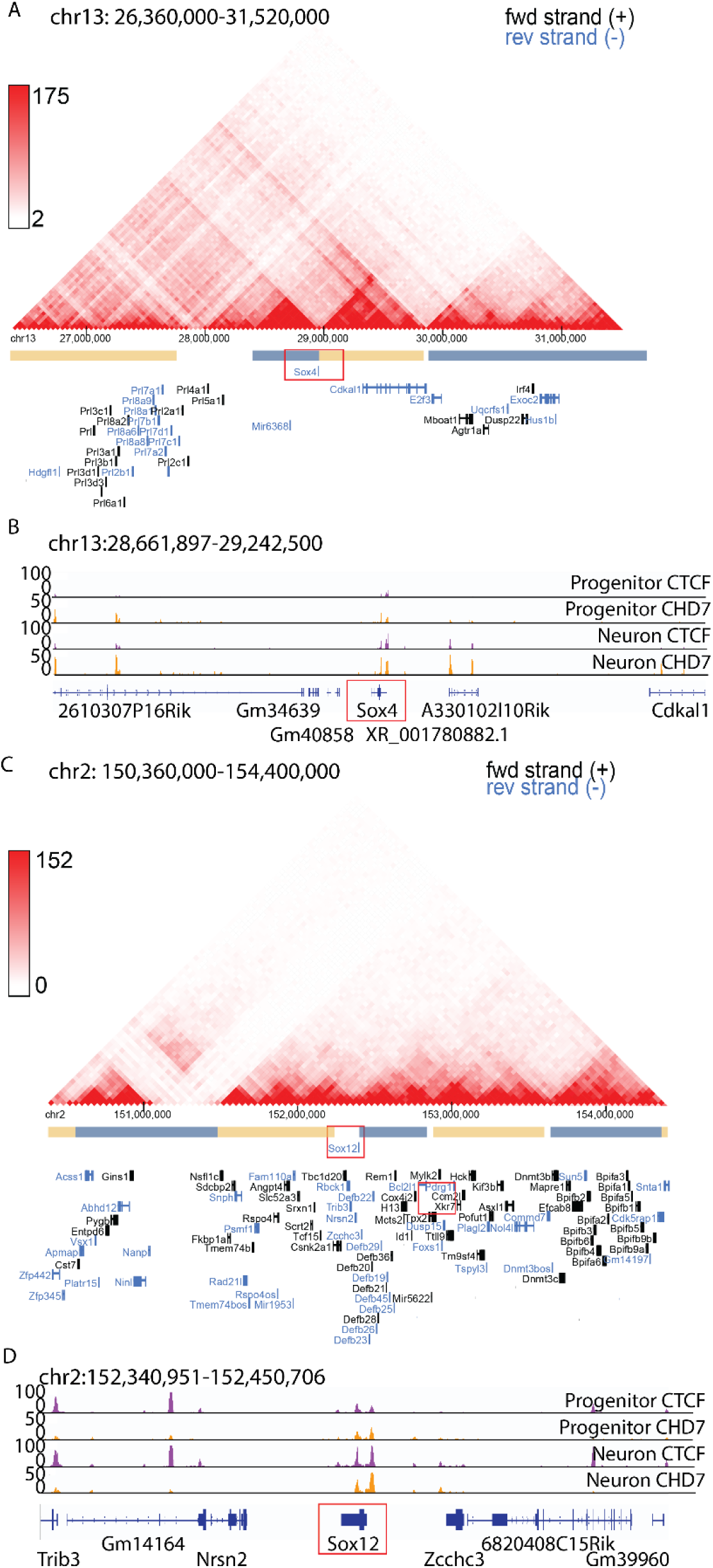
*SoxC* family of genes are located at TAD boundaries and display *Chd7* and CTCF marks. The SoxC family of transcription factors include *Sox4*, *Sox11* and *Sox12*. Contact maps for topologically associating domains (TADs) obtained from cortical neurons (Bonev et al., 2017) were visualized using the 3D Genome Browser (Wang et al., 2018). (A) Predicted TAD regions are indicated by blue and yellow bars below the contact map. *Sox4* resides at a boundary region between a pair of TADs. (B) Genome tracks around *Sox12* displayed CTCF+ CHD7+ peaks in both progenitors and neurons. (C) Similarly, *Sox12* is located at a TAD boundary, and (D) the genome tracks around *Sox12* show CTCF+ CHD7+ peaks in progenitors and neurons. Genomic coordinates for TADs and genomic tracks as labeled.

## References

Ahmed, M., Moon, R., Prajapati, R.S., James, E., Basson, M.A., and Streit, A. (2021). The chromatin remodelling factor Chd7 protects auditory neurons and sensory hair cells from stress-induced degeneration. Commun Biol 4, 1260.

Allis, C.D., and Jenuwein, T. (2016). The molecular hallmarks of epigenetic control. Nat Rev Genet 17, 487–500.

Benayoun, B.A., Pollina, E.A., Ucar, D., Mahmoudi, S., Karra, K., Wong, E.D., Devarajan, K., Daugherty, A.C., Kundaje, A.B., Mancini, E., et al. (2014). H3K4me3 breadth is linked to cell identity and transcriptional consistency. Cell 158, 673–688.

Bergsland, M., Werme, M., Malewicz, M., Perlmann, T., and Muhr, J. (2006). The establishment of neuronal properties is controlled by Sox4 and Sox11. Genes Dev 20, 3475–3486.

Bermingham, N.A., Hassan, B.A., Price, S.D., Vollrath, M.A., Ben-Arie, N., Eatock, R.A., Bellen, H.J., Lysakowski, A., and Zoghbi, H.Y. (1999). Math1: an essential gene for the generation of inner ear hair cells. Science 284, 1837–1841.

Bonev, B., Mendelson Cohen, N., Szabo, Q., Fritsch, L., Papadopoulos, G.L., Lubling, Y., Xu, X., Lv, X., Hugnot, J.P., Tanay, A., et al. (2017). Multiscale 3D Genome Rewiring during Mouse Neural Development. Cell 171, 557–572 e524.

Bouazoune, K., and Kingston, R.E. (2012). Chromatin remodeling by the CHD7 protein is impaired by mutations that cause human developmental disorders. Proc Natl Acad Sci U S A 109, 19238–19243.

Clapier, C.R., Iwasa, J., Cairns, B.R., and Peterson, C.L. (2017). Mechanisms of action and regulation of ATP-dependent chromatin-remodelling complexes. Nat Rev Mol Cell Biol 18, 407–422.

Dixon, J.R., Selvaraj, S., Yue, F., Kim, A., Li, Y., Shen, Y., Hu, M., Liu, J.S., and Ren, B. (2012). Topological domains in mammalian genomes identified by analysis of chromatin interactions. Nature 485, 376–380.

Doench, J.G., Fusi, N., Sullender, M., Hegde, M., Vaimberg, E.W., Donovan, K.F., Smith, I., Tothova, Z., Wilen, C., Orchard, R., et al. (2016). Optimized sgRNA design to maximize activity and minimize off-target effects of CRISPR-Cas9. Nat Biotechnol 34, 184–191.

Elkon, R., Milon, B., Morrison, L., Shah, M., Vijayakumar, S., Racherla, M., Leitch, C.C., Silipino, L., Hadi, S., Weiss-Gayet, M., et al. (2015). RFX transcription factors are essential for hearing in mice. Nat Commun 6, 8549.

Evsen, L., Sugahara, S., Uchikawa, M., Kondoh, H., and Wu, D.K. (2013). Progression of neurogenesis in the inner ear requires inhibition of Sox2 transcription by neurogenin1 and neurod1. J Neurosci 33, 3879–3890.

Finocchiaro, G., Carro, M.S., Francois, S., Parise, P., DiNinni, V., and Muller, H. (2007). Localizing hotspots of antisense transcription. Nucleic Acids Res 35, 1488–1500.

Flores, E.N., Duggan, A., Madathany, T., Hogan, A.K., Marquez, F.G., Kumar, G., Seal, R.P., Edwards, R.H., Liberman, M.C., and Garcia-Anoveros, J. (2015). A non-canonical pathway from cochlea to brain signals tissue-damaging noise. Curr Biol 25, 606–612.

Gao, J., Skidmore, J.M., Cimerman, J., Ritter, K.E., Qiu, J., Wilson, L.M.Q., Raphael, Y., Kwan, K.Y., and Martin, D.M. (2024). CHD7 and SOX2 act in a common gene regulatory network during mammalian semicircular canal and cochlear development. Proc Natl Acad Sci U S A 121, e2311720121.

Gerdes, J., Lemke, H., Baisch, H., Wacker, H.H., Schwab, U., and Stein, H. (1984). Cell cycle analysis of a cell proliferation-associated human nuclear antigen defined by the monoclonal antibody Ki-67. J Immunol 133, 1710–1715.

Gerdes, J., Schwab, U., Lemke, H., and Stein, H. (1983). Production of a mouse monoclonal antibody reactive with a human nuclear antigen associated with cell proliferation. Int J Cancer 31, 13–20.

Gnedeva, K., and Hudspeth, A.J. (2015). SoxC transcription factors are essential for the development of the inner ear. Proc Natl Acad Sci U S A 112, 14066–14071.

Grandi, F.C., De Tomasi, L., and Mustapha, M. (2020). Single-Cell RNA Analysis of Type I Spiral Ganglion Neurons Reveals a Lmx1a Population in the Cochlea. Front Mol Neurosci 13, 83.

Gu, Z., and Hubschmann, D. (2023). rGREAT: an R/bioconductor package for functional enrichment on genomic regions. Bioinformatics 39.

Guttman, M., Amit, I., Garber, M., French, C., Lin, M.F., Feldser, D., Huarte, M., Zuk, O., Carey, B.W., Cassady, J.P., et al. (2009). Chromatin signature reveals over a thousand highly conserved large non-coding RNAs in mammals. Nature 458, 223–227.

He, Y., Vogelstein, B., Velculescu, V.E., Papadopoulos, N., and Kinzler, K.W. (2008). The antisense transcriptomes of human cells. Science 322, 1855–1857.

Huang, H., Zhu, Q., Jussila, A., Han, Y., Bintu, B., Kern, C., Conte, M., Zhang, Y., Bianco, S., Chiariello, A.M., et al. (2021). CTCF mediates dosage- and sequence-context-dependent transcriptional insulation by forming local chromatin domains. Nat Genet 53, 1064–1074.

Hurd, E.A., Adams, M.E., Layman, W.S., Swiderski, D.L., Beyer, L.A., Halsey, K.E., Benson, J.M., Gong, T.W., Dolan, D.F., Raphael, Y., et al. (2011). Mature middle and inner ears express Chd7 and exhibit distinctive pathologies in a mouse model of CHARGE syndrome. Hear Res 282, 184–195.

Hurd, E.A., Capers, P.L., Blauwkamp, M.N., Adams, M.E., Raphael, Y., Poucher, H.K., and Martin, D.M. (2007). Loss of Chd7 function in gene-trapped reporter mice is embryonic lethal and associated with severe defects in multiple developing tissues. Mamm Genome 18, 94–104.

Hurd, E.A., Poucher, H.K., Cheng, K., Raphael, Y., and Martin, D.M. (2010). The ATP-dependent chromatin remodeling enzyme CHD7 regulates pro-neural gene expression and neurogenesis in the inner ear. Development 137, 3139–3150.

Huynh-Thu, V.A., Irrthum, A., Wehenkel, L., and Geurts, P. (2010). Inferring regulatory networks from expression data using tree-based methods. PLoS One 5.

Jadali, A., Song, Z., Laureano, A.S., Toro-Ramos, A., and Kwan, K. (2016). Initiating Differentiation in Immortalized Multipotent Otic Progenitor Cells. J Vis Exp.

Jahan, I., Kersigo, J., Pan, N., and Fritzsch, B. (2010). Neurod1 regulates survival and formation of connections in mouse ear and brain. Cell Tissue Res 341, 95–110.

Jahan, I., Pan, N., Kersigo, J., and Fritzsch, B. (2013). Beyond generalized hair cells: molecular cues for hair cell types. Hear Res 297, 30–41.

Kent, W.J., Zweig, A.S., Barber, G., Hinrichs, A.S., and Karolchik, D. (2010). BigWig and BigBed: enabling browsing of large distributed datasets. Bioinformatics 26, 2204–2207.

Kim, D., Langmead, B., and Salzberg, S.L. (2015). HISAT: a fast spliced aligner with low memory requirements. Nat Methods 12, 357–360.

Koundakjian, E.J., Appler, J.L., and Goodrich, L.V. (2007). Auditory neurons make stereotyped wiring decisions before maturation of their targets. J Neurosci 27, 14078–14088.

Kutner, R.H., Zhang, X.Y., and Reiser, J. (2009). Production, concentration and titration of pseudotyped HIV-1-based lentiviral vectors. Nature protocols 4, 495–505.

Kwan, K.Y., Shen, J., and Corey, D.P. (2015). C-MYC transcriptionally amplifies SOX2 target genes to regulate self-renewal in multipotent otic progenitor cells. Stem Cell Reports 4, 47–60.

Langmead, B., and Salzberg, S.L. (2012). Fast gapped-read alignment with Bowtie 2. Nat Methods 9, 357–359.

Li, C., Li, X., Bi, Z., Sugino, K., Wang, G., Zhu, T., and Liu, Z. (2020). Comprehensive transcriptome analysis of cochlear spiral ganglion neurons at multiple ages. Elife 9.

Li, H., Handsaker, B., Wysoker, A., Fennell, T., Ruan, J., Homer, N., Marth, G., Abecasis, G., Durbin, R., and Genome Project Data Processing, S. (2009). The Sequence Alignment/Map format and SAMtools. Bioinformatics 25, 2078–2079.

Ling, K.H., Hewitt, C.A., Beissbarth, T., Hyde, L., Banerjee, K., Cheah, P.S., Cannon, P.Z., Hahn, C.N., Thomas, P.Q., Smyth, G.K., et al. (2009). Molecular networks involved in mouse cerebral corticogenesis and spatio-temporal regulation of Sox4 and Sox11 novel antisense transcripts revealed by transcriptome profiling. Genome Biol 10, R104.

Liu, C., Glowatzki, E., and Fuchs, P.A. (2015). Unmyelinated type II afferent neurons report cochlear damage. Proc Natl Acad Sci U S A 112, 14723–14727.

Liu, M., Pereira, F.A., Price, S.D., Chu, M.J., Shope, C., Himes, D., Eatock, R.A., Brownell, W.E., Lysakowski, A., and Tsai, M.J. (2000). Essential role of BETA2/NeuroD1 in development of the vestibular and auditory systems. Genes Dev 14, 2839–2854.

Love, M.I., Huber, W., and Anders, S. (2014). Moderated estimation of fold change and dispersion for RNA-seq data with DESeq2. Genome Biol 15, 550.

Meers, M.P., Tenenbaum, D., and Henikoff, S. (2019). Peak calling by Sparse Enrichment Analysis for CUT&RUN chromatin profiling. Epigenetics Chromatin 12, 42.

Micucci, J.A., Sperry, E.D., and Martin, D.M. (2015). Chromodomain helicase DNA-binding proteins in stem cells and human developmental diseases. Stem Cells Dev 24, 917–926.

Narita, T., Ito, S., Higashijima, Y., Chu, W.K., Neumann, K., Walter, J., Satpathy, S., Liebner, T., Hamilton, W.B., Maskey, E., et al. (2021). Enhancers are activated by p300/CBP activity-dependent PIC assembly, RNAPII recruitment, and pause release. Mol Cell 81, 2166–2182 e2166.

Nie, J., Ueda, Y., Solivais, A.J., and Hashino, E. (2022). CHD7 regulates otic lineage specification and hair cell differentiation in human inner ear organoids. Nat Commun 13, 7053.

Nitarska, J., Smith, J.G., Sherlock, W.T., Hillege, M.M., Nott, A., Barshop, W.D., Vashisht, A.A., Wohlschlegel, J.A., Mitter, R., and Riccio, A. (2016). A Functional Switch of NuRD Chromatin Remodeling Complex Subunits Regulates Mouse Cortical Development. Cell Rep 17, 1683–1698.

Nitiss, J.L. (1998). Investigating the biological functions of DNA topoisomerases in eukaryotic cells. Biochim Biophys Acta 1400, 63–81.

Petitpre, C., Faure, L., Uhl, P., Fontanet, P., Filova, I., Pavlinkova, G., Adameyko, I., Hadjab, S., and Lallemend, F. (2022). Single-cell RNA-sequencing analysis of the developing mouse inner ear identifies molecular logic of auditory neuron diversification. Nat Commun 13, 3878.

Ramirez, F., Dundar, F., Diehl, S., Gruning, B.A., and Manke, T. (2014). deepTools: a flexible platform for exploring deep-sequencing data. Nucleic Acids Res 42, W187–191.

Rao, S.S., Huntley, M.H., Durand, N.C., Stamenova, E.K., Bochkov, I.D., Robinson, J.T., Sanborn, A.L., Machol, I., Omer, A.D., Lander, E.S., et al. (2014). A 3D map of the human genome at kilobase resolution reveals principles of chromatin looping. Cell 159, 1665–1680.

Reddy, N.C., Majidi, S.P., Kong, L., Nemera, M., Ferguson, C.J., Moore, M., Goncalves, T.M., Liu, H.K., Fitzpatrick, J.A.J., Zhao, G., et al. (2021). CHARGE syndrome protein CHD7 regulates epigenomic activation of enhancers in granule cell precursors and gyrification of the cerebellum. Nat Commun 12, 5702.

Robinson, J.T., Thorvaldsdottir, H., Winckler, W., Guttman, M., Lander, E.S., Getz, G., and Mesirov, J.P. (2011). Integrative genomics viewer. Nat Biotechnol 29, 24–26.

Sanders, T.R., and Kelley, M.W. (2022). Specification of neuronal subtypes in the spiral ganglion begins prior to birth in the mouse. Proc Natl Acad Sci U S A 119, e2203935119.

Sanosaka, T., Okuno, H., Mizota, N., Andoh-Noda, T., Sato, M., Tomooka, R., Banno, S., Kohyama, J., and Okano, H. (2022). Chromatin remodeler CHD7 targets active enhancer region to regulate cell type-specific gene expression in human neural crest cells. Sci Rep 12, 22648.

Sanson, K.R., Hanna, R.E., Hegde, M., Donovan, K.F., Strand, C., Sullender, M.E., Vaimberg, E.W., Goodale, A., Root, D.E., Piccioni, F., et al. (2018). Optimized libraries for CRISPR-Cas9 genetic screens with multiple modalities. Nat Commun 9, 5416.

Schnetz, M.P., Bartels, C.F., Shastri, K., Balasubramanian, D., Zentner, G.E., Balaji, R., Zhang, X., Song, L., Wang, Z., Laframboise, T., et al. (2009). Genomic distribution of CHD7 on chromatin tracks H3K4 methylation patterns. Genome Res 19, 590–601.

Shrestha, B.R., Chia, C., Wu, L., Kujawa, S.G., Liberman, M.C., and Goodrich, L.V. (2018). Sensory Neuron Diversity in the Inner Ear Is Shaped by Activity. Cell 174, 1229–1246 e1217.

Song, Z., Jadali, A., Fritzsch, B., and Kwan, K.Y. (2017). NEUROG1 Regulates CDK2 to Promote Proliferation in Otic Progenitors. Stem Cell Reports 9, 1516–1529.

Su, Y.X., Hou, C.C., and Yang, W.X. (2015). Control of hair cell development by molecular pathways involving Atoh1, Hes1 and Hes5. Gene 558, 6–24.

Sun, M., Hurst, L.D., Carmichael, G.G., and Chen, J. (2005). Evidence for a preferential targeting of 3’-UTRs by cis-encoded natural antisense transcripts. Nucleic Acids Res 33, 5533–5543.

Sun, Y., Wang, L., Zhu, T., Wu, B., Wang, G., Luo, Z., Li, C., Wei, W., and Liu, Z. (2022). Single-cell transcriptomic landscapes of the otic neuronal lineage at multiple early embryonic ages. Cell Rep 38, 110542.

Sun, Z., Zhang, Y., Jia, J., Fang, Y., Tang, Y., Wu, H., and Fang, D. (2020). H3K36me3, message from chromatin to DNA damage repair. Cell Biosci 10, 9.

Torrado, M., Low, J.K.K., Silva, A.P.G., Schmidberger, J.W., Sana, M., Sharifi Tabar, M., Isilak, M.E., Winning, C.S., Kwong, C., Bedward, M.J., et al. (2017). Refinement of the subunit interaction network within the nucleosome remodelling and deacetylase (NuRD) complex. FEBS J 284, 4216–4232.

van Ravenswaaij-Arts, C., and Martin, D.M. (2017). New insights and advances in CHARGE syndrome: Diagnosis, etiologies, treatments, and research discoveries. Am J Med Genet C Semin Med Genet 175, 397–406.

Wang, Y., Song, F., Zhang, B., Zhang, L., Xu, J., Kuang, D., Li, D., Choudhary, M.N.K., Li, Y., Hu, M., et al. (2018). The 3D Genome Browser: a web-based browser for visualizing 3D genome organization and long-range chromatin interactions. Genome Biol 19, 151.

Wegner, M. (2011). SOX after SOX: SOXession regulates neurogenesis. Genes Dev 25, 2423–2428.

Xu, M., Li, S., Xie, X., Guo, L., Yu, D., Zhuo, J., Lin, J., Kol, L., and Gan, L. (2024). ISL1 and POU4F1 Directly Interact to Regulate the Differentiation and Survival of Inner Ear Sensory Neurons. J Neurosci 44.

Yeo, N.C., Chavez, A., Lance-Byrne, A., Chan, Y., Menn, D., Milanova, D., Kuo, C.C., Guo, X., Sharma, S., Tung, A., et al. (2018). An enhanced CRISPR repressor for targeted mammalian gene regulation. Nat Methods 15, 611–616.

Zhang, Y., Liu, T., Meyer, C.A., Eeckhoute, J., Johnson, D.S., Bernstein, B.E., Nusbaum, C., Myers, R.M., Brown, M., Li, W., et al. (2008). Model-based analysis of ChIP-Seq (MACS). Genome Biol 9, R137.

Zhang, Y., Liu, X.S., Liu, Q.R., and Wei, L. (2006). Genome-wide in silico identification and analysis of cis natural antisense transcripts (cis-NATs) in ten species. Nucleic Acids Res 34, 3465–3475.

Zine, A., Aubert, A., Qiu, J., Therianos, S., Guillemot, F., Kageyama, R., and de Ribaupierre, F. (2001). Hes1 and Hes5 activities are required for the normal development of the hair cells in the mammalian inner ear. J Neurosci 21, 4712–4720.

